# The Single-Cell Transcriptome Program of Nodule Development Cellular Lineages in *Medicago truncatula*

**DOI:** 10.1101/2023.06.13.544787

**Authors:** Wendell J. Pereira, Jade Boyd, Daniel Conde, Paolo M. Triozzi, Kelly M. Balmant, Christopher Dervinis, Henry W. Schmidt, Carolina Boaventura-Novaes, Sanhita Chakraborty, Sara A. Knaack, Yueyao Gao, Frank Alexander Feltus, Sushmita Roy, Jean-Michel Ané, Julia Frugoli, Matias Kirst

## Abstract

Legumes can establish a symbiotic relationship with nitrogen-fixing rhizobia by developing nodules after root exposure to lipo-chito-oligosaccharides secreted by the bacteria. Nodule development initiates with anticlinal mitotic divisions in the pericycle and endodermal and inner cortical cells, establishing cell lineages that ultimately form each nodule compartment. We characterized these lineages by isolating and sequencing the transcriptome of *Medicago truncatula* single nuclei derived from uninoculated roots and roots undergoing early nodule development at 24, 48, and 96 hours after inoculation. To enrich samples for cells responding to the rhizobia, we complemented the analysis of the *Medicago* wild-type genotype A17 with a mutant for the autoregulation of nodulation, *sunn-4*. Analysis of cell lineage trajectories derived from the cortex indicates that their transcriptome is initially enriched for cytokinin perception and signaling while repressing auxin accumulation. As these cells differentiate to form nodules, expression of genes related to auxin biosynthesis, transport, and signaling was enhanced, while genes involved in cytokinin degradation were activated as lineages bifurcated to form the nodule meristem and infection zones. While the contribution of auxin and cytokinin in nodule development has been recognized, this single-cell resource quantifies the expression of each of their regulators, receptors and targets as cells divide and differentiate to form each nodule compartment.

## INTRODUCTION

Nitrogen is an essential nutrient for plant growth. To acquire nitrogen, plants of the legume family establish a symbiosis with rhizobia bacteria that leads to the development of nodules on legume roots. These nodules are *de novo* plant organs that provide an optimal environment for the rhizobia to fix nitrogen in exchange for photosynthates. The establishment of root nodule symbioses (RNS) requires the coordination of two distinct processes: bacterial infection and nodule organogenesis. In the model legume *Medicago truncatula*, in response to the rhizobium *Sinorhizobium meliloti*, the nucleus of the root hair cells increases in size, and genes involved in defense responses are transiently activated, together with bacterial lipo-chito-oligosaccharides (LCOs) signaling within the first 24 hours of interaction (Breakspear *et al*., 2014b; Timmers *et al*., 1999). In parallel, cell divisions occur in the pericycle to initiate nodule organogenesis. This first mitotic event is followed by the division of cortex cells that differentiate into the nodule primordium (Timmers *et al*., 1999; Xiao *et al*., 2014). Together with the endodermis, the pericycle-derived cells differentiate into the uninfected cells at the base of nodules, while inner-cortex cells originate the other nodule compartments.

Nodules are modified lateral roots (Hirsch *et al*., 1997; Schiessl *et al*., 2019). Their development is primarily regulated by the interplay between the phytohormones auxin and cytokinin (CK), synthesized and transported across specific root cells (Lin *et al*., 2020). Rhizobia induce the biosynthesis of CK by *LONELY GUY 3* (*LOG3*) in the epidermis (Jardinaud *et al*., 2016), which is translocated to inner root cells by the ABCG CK transporter *ABCG56* (Jarzyniak *et al*., 2021). Cytokinin signaling in the root cortex and pericycle activates the expression of transcription factors such as *NODULE INCEPTION* (*NIN*) and the biosynthesis and accumulation of auxin through the induction of *LOB-DOMAIN PROTEIN 16* (*LBD16*) by *NIN* (Schiessl *et al*., 2019). Because of the biological cost to the plant, the host tightly regulates nodule development through the autoregulation of nodulation (AON, reviewed by Ferguson *et al*., 2019). After rhizobia infection, peptides of the CLE (CLAVATA3 (CL3)/endosperm-surrounding region) family are induced and transported to the shoot. These peptides are perceived by a leucine-rich repeat receptor-like kinase (LRR-RLK) in the shoot, triggering the AON pathway by releasing a mobile signal. The LRR-RLK is encoded by *SUNN* (*SUPER NUMERIC NODULES*) in *M. truncatula* (Schnabel *et al*., 2005). A null mutation in *SUNN* (*sunn-4*) results in a significant increase in the number of nodules (>5-fold) (Schnabel *et al*., 2010).

The molecular mechanisms involved in the activation or repression of gene expression during nodule development have been extensively examined by transcriptome analysis of whole roots from nodulating plants (reviewed by Mergaert *et al*., 2020). In these studies, the response of each cell is masked by the average bulk RNA profile, a critical limitation considering the importance of distinct root cell types assuming specific roles in nodule development (Xiao *et al*., 2014). Recently developed approaches such as single-cell RNA sequencing (scRNA-seq) allow for uncovering cell transcriptional heterogeneity within a tissue or organ (Giacomello, 2021).

Furthermore, the application of novel analytical streategies, such as pseudotime analysis, adds a developmental and temporal dimension to single-cell genomics data, allowing for cell lineages to be inferred and genes that control the process to be identified (Saelens *et al*., 2019). Consequently, putative regulators of developmental processes such as shoot apex (Zhang *et al*., 2021; Conde *et al*., 2022), leaf (Liu *et al*., 2020; Lopez-Anido *et al*., 2021), and wood differentiation (Chen *et al*., 2021; Xie *et al*., 2022) have been determined and, in some cases, validated. Similar strategies could uncover the molecular regulators involved in the differentiation of root cell types as they transition into the components of a nodule. This requires data to be collected across the developmental process, a limitation of existing studies (Cervantes-Pérez *et al*., 2022; Ye *et al*., 2022).

Cells responding to rhizobia represent a small fraction of the root tissue, resulting in limitations for single-cell genomic analysis of nodule development, as few cells responding to the stimulus are present in the whole-root cell population. Here we present a high-resolution and cell type-specific gene expression map of the roots of the legume *M. truncatula* during early nodule development. To capture a significant fraction of cells responding to the rhizobia and strengthen the inferences about the molecular mechanisms involved in regulating nodulation, we complemented the wild-type genotype A17 (WT) analysis with data collected from the *sunn-4* mutant (Penmetsa *et al*., 2003), a regulator of the AON pathway. By disrupting the AON, the developmental program of nodules remains unaffected while producing a response to infection and triggering nodule development that affects an extensive fraction of the root cortex and pericycle cells (Penmetsa *et al*., 2003). Using this approach, we identified the infected root hair cells and characterized their transcriptional differences from the uninfected cells. In addition, we reconstructed the developmental trajectory as cortical cells differentiate to generate the nodule meristem and the infected cells of the mature nodule, identifying genes that govern this process and may be required for engineering nodule organogenesis and N-fixation in crop plants.

## RESULTS

To identify the distinct cell populations present in the sampled tissue, we performed clustering after combining the single-cell transcriptomes obtained in four time points (0 h, 24 hpi, 48 hpi, and 96 hpi) for each of the two genotypes (Jemalong A17 and *sunn-4*), individually, using Monocle 3 with default parameters. The combined datasets of each genotype were corrected for batch effects (see Methods) and comprised 18,635 nuclei grouped in 16 clusters for the WT (Figure S1-A) and 18,233 nuclei in 17 clusters for the *sunn-4* (Figure S1-B). The presence of similar cell types in both genotypes was confirmed by observing the expression of marker genes previously characterized in roots engaging in RNS (Figure S1-C). Next, to allow the comparison among genotypes, we combined all datasets (the four time-points and two genotypes) in a single analysis and identified 29 clusters (Figure 1A) containing 36,868 nuclei and 36,395 expressed genes. A web application has been developed for exploring the expression profile of *M. truncatula* genes during nodule development https://.

**Figure 1.**
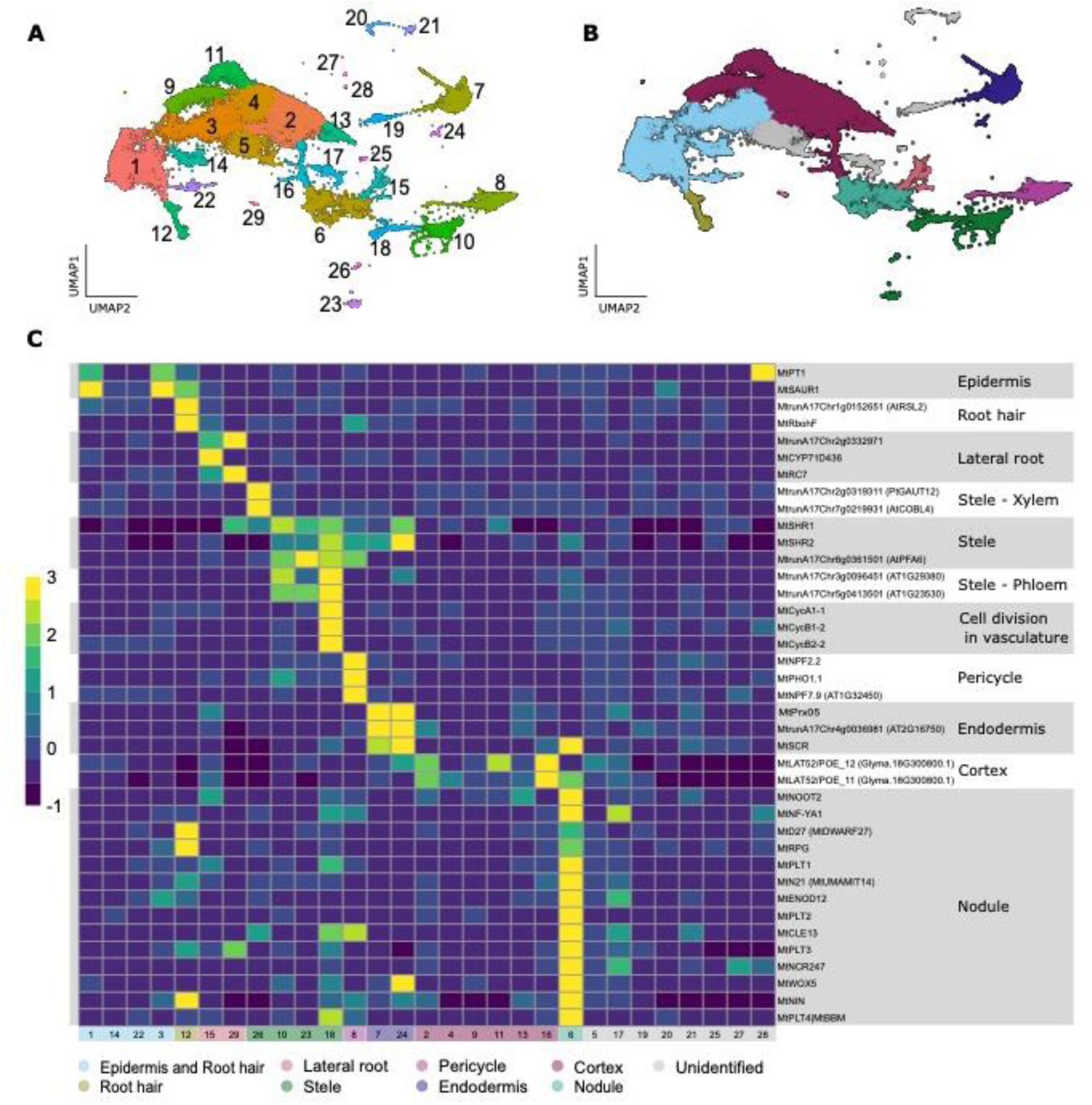
Single-nuclei transcriptomes from *M. truncatula* roots cluster into distinct cell types. **(A)** UMAP projection plot showing the distribution of the 36,868 individual cells from the two genotypes and four time points. Cells were grouped in 29 clusters according to their gene expression profiles. **(B)** Six cell types were identified, in addition to clusters containing cells from the lateral root and nodule development. **(C)** The expression of known cell-type marker genes across cells reveals the identity of clusters. In **(C)**, color shows the average expression across cells in each cluster. Additional file S1 contains the complete list of cell-type markers used in the present study.

To attribute cell identities to each cluster, we analyzed the expression profile of *M. truncatula* homologs of cell-type-specific markers previously identified in *Arabidopsis* roots (Denyer *et al*., 2019; Ryu *et al*., 2019; T.,-Q., Zhang *et al*., 2019; Shulse *et al*., 2019) (Additional file 1), and markers validated in *M. truncatula* or other species capable of RNS (Additional file 1). In addition, we evaluated the expression of genes specific to each cluster (Moran’s I test, q-value < 0.05) in a laser capture microdissection (LCM, Schnabel *et al*., 2023) dataset that investigated the response of *M. truncatula* to rhizobia infection at similar time points as the ones used in the present study (see Methods; Figures S2-S7). Based on the annotation approach described above, we identified nuclei derived from six types of root cells: the epidermal cells (clusters 1, 3, 14, and 22), root hair (clusters 12), cortex (clusters 2, 4, 9, 11, 13, and 16), endodermis (clusters 7 and 24), pericycle (cluster 8) and stele (excluding pericycle; clusters 10, 18, 23 and 26) (Figure 1B-C).

While the cluster annotation relied on a combination of strategies, some of the most informative marker genes for each cell type are described below. Epidermal cells were detected based on the expression of the phosphate transporter *MtPT1* (Damiani *et al*., 2016) and a homolog of the *Arabidopsis SMALL AUXIN UPREGULATED RNA* (*SAUR1*). Root hair cells were identified by the expression of *RESPIRATORY BURST OXIDASE HOMOLOG F* (*MtRbohF*, Marino *et al*., 2011). In cluster 7, the specific expression of *SCARECROW* (Roux *et al*., 2014) indicated it is composed of endodermal cells. The detection of cortical cells was supported by the expression of homologs of the *Glycine max* gene *Glyma.18G300800*, previously shown to be specifically expressed in this cell type (Song *et al*., 2022). The stele was characterized by the expression of *SHORT ROOT 1* and *SHORT ROOT 2* (Roux *et al*., 2014; Dong *et al*., 2021), besides markers specific for the xylem, including the homologs of Arabidopsis *GAUT12* (Persson *et al*., 2007) and *COBL4* (Lasserre *et al*., 2008), and the homologs of Arabidopsis *AT1G29380* and *AT1G23530*, uncharacterized genes described as markers of the phloem (T.,-Q., Zhang *et al*., 2019). Markers for early cell proliferation of the vascular procambial/cambial cells in the stele (*e.g.*, homologs of *Arabidopsis CYCLIN A1;1,* J., Zhang *et al*., 2019) were detected in a cluster related to cell division (cluster 18). In contrast, cluster 8 is highly distinct transcriptionally and characterized by the expression of markers for the pericycle, including homologs of *Arabidopsis NITRATE TRANSPORTER 1.5* genes (Lin *et al*., 2008). Notably, the annotation of all mentioned clusters was also supported by the expression profile of their most specific genes (100 most significant genes, q-value < 0.05, sorted by the specificity value for Moran’s I test) in the LCM data (Figures S2-S7) and additional marker genes listed in Additional file 1.

Clusters 15 and 29 were identified as representing cells from lateral roots based on the detection of genes expressed specifically during that process and unrelated to RNS (Schiessl *et al*., 2019) (Figure 1C, Figure S8-A). For eight clusters, it was not possible to identify their corresponding type, due to the lack of expression of known marker genes and absence of evidence pointing to a specific cell type in the genes specifically expressed in those clusters. Those include clusters 5 (n = 2,410 cells), 17 (n = 597), 19 (n = 414), 20 (n = 328), 21 (n = 321), 25 (n = 92), 27 (n = 62), and 28 (n = 38).

We also identified a cluster of cells involved in nodule development (cluster 6), as shown by the expression of essential genes related to RNS and of genes specifically expressed in that cluster in the LCM data (Figure 1C, Figure S8-B). Moreover, while cluster 16 is composed of cells from the cortex, the expression of its marker genes strongly overlaps with those detected in nodules in the LCM data, suggesting that this cluster represents cells transitioning from cortex to nodule.

### Changes in the function of cell types after infection

After clustering and annotating the cells detected throughout the experiment (0, 24, 48, and 96 hpi) in both genotypes (wild-type A17 and the mutant *sunn-4*), we assessed the changes in gene expression and the emergence of novel cell types reflecting the root response to rhizobia. Additional file 2 describes the list of genes significantly enriched for expression in each cluster.

#### Root hair cells responding to infection

RNS begins with the perception of bacterial LCOs by root hair cells, followed by infection and the formation of an infection thread in these cells. As expected, at 24 hpi, the transcriptome profile of a subset of root hair cells (cluster 12) transitioned to express marker genes for the infection thread and infectosome, including *RHIZOBIUM DIRECTED POLAR GROWTH* (*MtRPG,* Arrighi *et al*., 2008; Figure S8-E), *CYSTATHIONINE-Β-SYNTHASE-LIKE DOMAIN-CONTAINING* (*MtCBS1*, Sinharoy *et al*., 2016; Figure S8-E), *NODULE PECTATE LYASE* (*MtNPL*, Xie *et al*., 2012), *CHALCONE O-METHYL TRANSFERASE* (*MtChOMT1*, Maxwell *et al*., 1993), *FLOTILLIN 4* (*MtFLOT4*, Liang *et al*., 2018) and *VAPYRIN* (*MtVPY*, Murray *et al*., 2011). Cluster 12 also showed the highest expression of genes related to epidermal infection (Schiessl *et al*., 2019) (Figure S8-E). Interestingly, we also detected a high expression of several of these genes in cluster 6, composed of cells from the developing nodule, pointing to their participation in the infection of the cortex-derived cells in the new root organ (Figure S8-E).

#### Pericycle cell division

The first cell divisions triggered by the rhizobia occur in the pericycle as early as 24 hpi (Xiao *et al*., 2014). We observed that a subset of the pericycle cells (cluster 8, Figure S9) differentiated transcriptionally from other cells in this cluster after infection. The appearance of these cells coincided with the expression of essential RNS genes in this cluster, such as *ISOPENTENYLTRANSFERASE 3* (*MtIPT3,* Triozzi *et al*., 2022), *CLAVATA3/EMBRYO-SURROUNDING REGION* 12 and 13 (*MtCLE12* and *MtCLE13*, Mortier *et al*., 2010), *MtENOD20* (Greene *et al*., 1998), *C-TERMINALLY ENCODED PEPTIDE1* (*MtCEP1*, de Bang *et al*., 2017), *NODULE INCEPTION* (*MtNIN,* Barnett *et al*., 2004) and *NUCLEAR FACTOR Y* (*MtNF-YA1*, Combier *et al*., 2006).

#### Early nodule development

Two clusters appeared during the experiment in A17 and *sunn-4*, reflecting the emergence of new cell types induced by the rhizobial infection. Cells in clusters 16 and 6 were first noted at 48 hpi, followed by an expansion at 96 hpi (Figures S9 and S10). Due to its hypernodulating phenotype that enriched the samples for cells from developing nodules, the increase in the number of cells in both clusters was more pronounced in the *sunn-4* genotype (Figure S10).

Cluster 16 comprised cortex cells responding to the infection, transitioning towards nodule formation (nodule primordia). This cluster did not strongly express marker genes associated with mature nodules (Figure S8-B). However, the transcription profile of its most specific markers strongly overlaps the expression detected in the LCM data obtained from developing nodules (Figure S5). Moreover, markers for the nodule primordia such as *NON SPECIFIC LIPID TRANSFER PROTEIN* (*MtN5* (Denny, 1996)), *BASIC HELIX-LOOP-HELIX 1* (*MtbHLH1*, Godiard *et al*., 2011), and the *AUXIN INFLUX TRANSPORTER LAX2* (*MtLAX2*, Roy *et al*., 2017), were strongly expressed in this cluster, reinforcing the hypothesis that these cell are in a developmentally transitioning state.

Cluster 6 was composed of cells that are part of the developing nodule, as shown by the localized expression of multiple nodule-specific genes (Schiessl *et al*., 2019; Figure S 8-B) and by the profile of the most specifically expressed genes of this cluster, in the LCM data (Figure S3). Based on the expression of known markers, cells from two of the nodule zones could be detected at 96 hpi as part of cluster 6. These include cells from the nodule meristem, expressing *PLETHORA* (*MtPLT*) 1, 3, and 4 (Franssen *et al*., 2015), *WUSCHEL-RELATED HOMEOBOX 5* (*MtWOX5*, Franssen *et al*., 2015), *MtCLE13* (Mortier *et al*., 2010) and *NODULE ROOT2* (*MtNOOT2*, Magne *et al*., 2018). Also detected were genes expressed in the infection zone (ZII), such as *NODULE-SPECIFIC PLAT-DOMAIN 1* (*MtNPD1*, Van de Velde *et al*., 2010), *MtNF-YA1* (Laporte *et al*., 2014), *NODULE-SPECIFIC CYSTEINE-RICH PEPTIDE 247* (*MtNCR247*, Farkas *et al*., 2014), *USUALLY MULTIPLE ACIDS MOVE IN AND OUT TRANSPORTERS 14* (*MtUMAMIT14*/*MtN21*, Garcia *et al*., 2023) and *SWEET11* (Kryvoruchko *et al*., 2016).

We searched for cells expressing NCR peptides to identify those representing the nitrogen-fixing zone (ZIII). The *M. truncatula* genome encodes for hundreds of such peptides, many of which have been associated with bacteroid differentiation (*e.g., MtNCR247*), acting in the cells where bacterial cell division is arrested, and cell elongation starts (Farkas *et al*., 2014). Investigating the expression profile of all 675 NCR peptides in version 5 of the *M. truncatula* genome revealed that genes encoding for 272 were expressed in our dataset (Additional file 1). A large fraction was expressed in the nodule cluster (cluster 6; Figure S11). However, NCR peptides specific to the nitrogen-fixing zone (ZIII), such as the *MtNCR035* and *MtNCR001* (Van de Velde *et al*., 2010), were not detected. The lack of transcripts for both peptides suggests an absence of cells from the ZIII, compatible with the developing stage of the nodules at 96 hpi in *M. truncatula* (Xiao *et al*., 2014). Alternatively, the number of cells from ZIII may be too limited to result in a discernible cluster.

### Higher resolution single-cell analysis of the root reveals cell type changes of the transcriptome triggered by RNS

Next, we separated cells from the complete dataset into two subsets related to distinct cellular responses to RNS. More specifically, cells associated with the response of root hair to the rhizobia and the formation of the infection thread; the inner-cortex cells of the nodule primordium and, later, differentiate into the meristem and the infected cells in the infection zone, were analyzed individually. Each of these subsets of cells was reclustered to identify the transcriptional changes triggered by RNS.

#### Rhizobia-infected root hair cells and formation of the infection thread

Rhizobia infect root hairs, resulting in the rapid establishment of infection threads in these cells. For this analysis, cells initially annotated as root hair and epidermis (Figure 1, clusters 1, 12, 14, and 22; total of 7,534 nuclei) were reclustered to identify those responding to the rhizobia within the first 24 hpi. Cluster 3, also annotated as the epidermis, was not included in this analysis because it contains a significant fraction of cortical cells (Figure S2).

Reanalysis of root hair and epidermal cells identified 12 clusters (Figure 2), hereinafter referred to as RH1 to RH12, of which RH10 was absent at 0 h but appeared at 24 hpi in the genotypes A17 and *sunn-4* (Figure S12-A). Annotation based on marker genes for root hair containing infection thread, including *MtFLOT4*, *MtVPY, MtRPG*, *MtCBS1*, *MtChOMT1*, and *MtNPL,* confirmed the identity of this emerging cluster. By investigating the expression of marker genes (*e.g.*, *MtRbohF*, Marino *et al*., 2011), and the list of most specifically expressed genes in each cluster (Additional file 2), we identified RH8 as representing root hair. Furthermore, the expression profile of genes previously identified as markers for epidermal infection (Schiessl *et al*., 2019) showed that RH8 and RH10 are involved in this process (Figure 2D).

**Figure 2.**
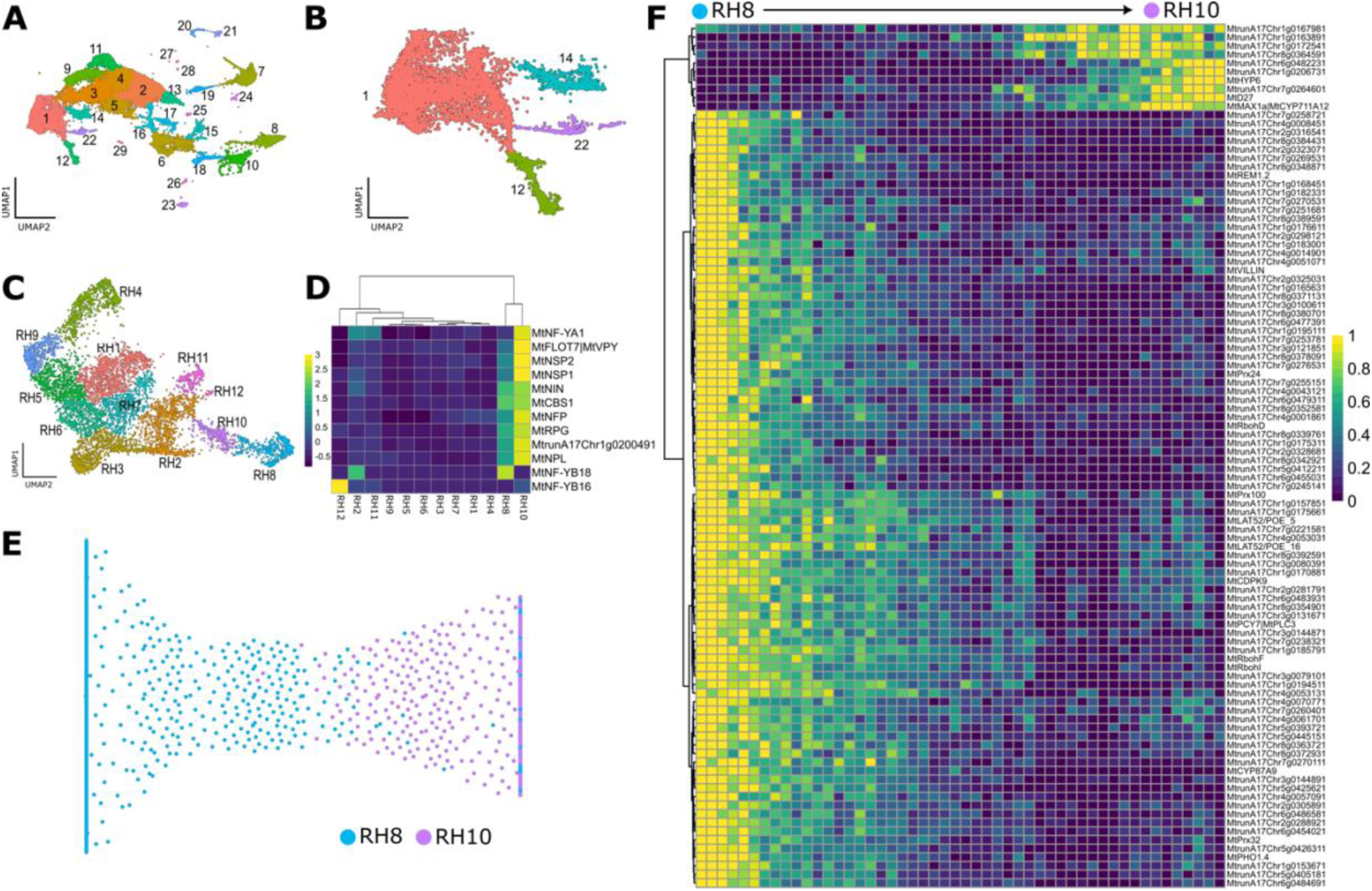
Analysis of epidermal and root hair cells during RNS identifies transcriptional changes triggered by infection. From the whole dataset **(A)**, cells belonging to clusters 1, 12, 14, and 22 were isolated **(B)** and reclustered in 12 clusters using a resolution of 0.001 **(C)**. Using marker genes and markers of epidermal infection identified previously (Schiessl *et al*., 2019), RH8 was identified as root hair and RH10 as root hair containing infection threads **(D)**. The trajectory inferred from RH8 **(E)**. The expression of the 100 most significant DEGs (TradeSeq, FDR < 0.001, sorted by the Wald test statistic) is shown along the trajectory **(F)** (Additional file 3). Cells in the trajectory were binned into 50 continuous groups of equal size to facilitate data visualization.

Next, we identified the genes most specifically expressed in the infected cells of RH10 (Moran’s I test, q-value < 0.05) and, therefore, differentially expressed compared to the remaining cells (Additional file 2). In addition to the specific genes, the visual comparison of the expression profiles in the UMAP plots showed that several LysM domain-containing receptor kinases are induced in infected root hair cells. In legumes, LCOs secreted by rhizobia are detected by a heteromeric Lys motif (LysM)-type transmembrane receptor complex. In *Medicago*, this association has been shown to involve the LYK-I and LYR-type receptors *LYSM DOMAIN CONTAINING RECEPTOR KINASE 3* (*MtLYK3*) and *NOD FACTOR PERCEPTION* (*MtNFP*), respectively (Limpens *et al*., 2003). Both of those genes are transcribed in cells of cluster RH10, besides being expressed in cells from other clusters. In addition, the *E3 UBIQUITIN LIGASE PUB1* (*MtPUB1|MtPUB45*), which has been shown to interact with *MtLYK3* (Mbengue *et al*., 2010), is specifically expressed in root hair-containing infection threads. Our data also indicate that genes encoding for other LysM domain-containing receptor kinases are expressed specifically in root hair-containing infection threads (*MtLYK4* and *MtLYK10*). Expectedly, *MtNFP* and *MtLYK3* were differentially expressed in the trajectory between uninfected and infected root hair cells (Additional file 3).

Reactive oxygen species (ROS) are abundant during nodulation and mainly generated by NADPH oxidases, or respiratory burst oxidase homologs (RBOHs). The Medicago RBOH gene family contains ten members, some of which catalyze the generation of ROS in response to infection when a protein complex is formed with calcium-dependent protein kinases (CDPK, Yu *et al*., 2018). When comparing the expression of RBOHs and CDPKs, we observed a significant, specific expression of *MtRbohB, MtRbohD, MtRbohF*, *MtRbohH, MtRbohI,* and *MtCDPK9* in root hair cells lacking infection threads (RH8; Moran’s test, q-value < 0.05). When considering differences in expression between uninfected (RH8) and infected root hair (RH10) cells within the trajectory, *MtRbohB*, *MtRbohD, MtRbohF*, *MtRbohI,* and *MtCDPK9* were significantly (TradeSeq, FDR < 0.001) more highly transcribed at the beginning of the trajectory (RH8, Figure 2), pointing to a coordinated activation of innate immunity in those cells, in contrast with those containing infection threads where their expression is suppressed.

Strigolactones (SL) are a class of phytohormones that promote infection thread formation (McAdam *et al*., 2017). Expectedly, the expression of genes in the SL biosynthesis pathway, such as *MtMAX1a* and *MtD27* (Mashiguchi *et al*., 2021), were among the most specific to RH10 (Moran’s test, q-value < 0.05, Additional file 2). Additionally, genes encoding for proteins involved in the synthesis of precursors of SL (carotenoids), including *MtZDS* and *MtZ-ISO*, were also identified as differentially regulated, with higher expression in infected root hair cells (RH10) compared to the remaining cells (Figure S13; Moran’s test, q-value < 0.05). In contrast, essential SL biosynthesis genes (*MtCCD7* and *MtCCD8*) previously found to be expressed specifically in infected root hair and developing nodule primordia (Breakspear *et al*., 2014a) were not detected as significantly differentially expressed. A visual inspection of their expression in the UMAP plot reveals a few cells expressing these genes, specifically in RH10 (Figure S13).

#### Cortex and nodule development

In response to rhizobium, legumes activate cytokinin signaling and local auxin accumulation in cortical root cells. The balance of these phytohormones regulates the mitotic division of distinct cortical cell layers, originating cell lineages that form each nodule compartment (Xiao *et al*., 2014). Because the abundance of cells from developing nodules was more pronounced in the hypernodulating *sunn-4* mutant (Figure S10, clusters 6 and 16), we assessed transcriptional changes during cortex differentiation in single-cell data derived from that genotype only. For the analysis, root cortex cells (Figure 1, cluster 2) detected at all time points (Figure S9) and clusters representing cell lineages that emerged from those cortex cells at 96 hpi (Figure 1, clusters 6 and 16), were evaluated. We noticed that a subset of cells from cluster 6 was already present at 0 hpi. Those cells were identified as lateral root meristem and removed from the dataset prior to reclustering (Figure S14). Reclustering resulted in the detection of eight groups of cells, representing 3,850 nuclei, hereinafter referred to as CN1 to CN8, the majority of which (CN3, CN4, CN5, CN6, CN7, and CN8) emerged at 96 hpi and are associated with nodule development (Figure 3 and Figure 4).

**Figure 3.**
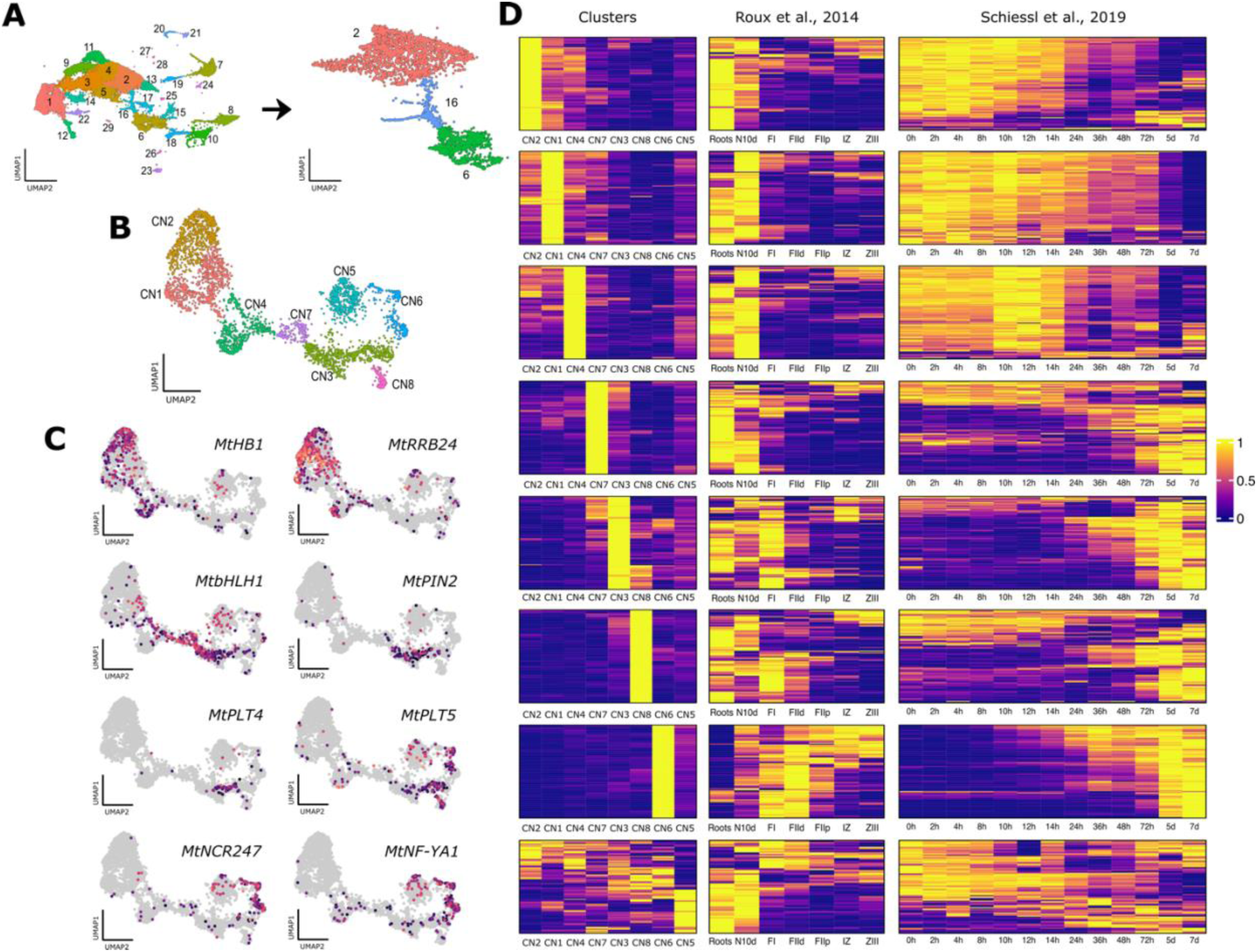
Reclustering of *sunn-4* cells in high resolution reveals distinct nodule components. **(A)** From the dataset obtained from the mutant *sunn-4,* cells belonging to clusters 2, 16, and 6 were isolated. **(B)** UMAP plot showing the distribution of cells in the eight clusters obtained after reclustering cells from cortex and nodules. **(C)** UMAP plots representing the expression profile of marker genes for cortex (*MtHB1* and *MtRRB24*), nodule primordium (*MtbHLH1* and *MtPIN2*), nodule meristem (*MtPLT4* and *MtPLT5*), and for infected cells from the infection zone (*MtNCR247* and *MtNF-YA1*). **(D)** Expression of the most specific genes (n=100; q-value < 0.05, sorted by the specificity value returned by the Moran’s I test) of each of the eight clusters (left panel) and their transcriptional profile in two reference datasets, including nodule fractions obtained by laser capture microdissection (Roux *et al*., 2014; middle panel) and bulk RNA sequencing (Schiessl *et al*., 2019; right panel). Each dataset was scaled and row-centered separately. The clusters are shown in the order they appear in the UMAP plots represented in **(A)**.

**Figure 4.**
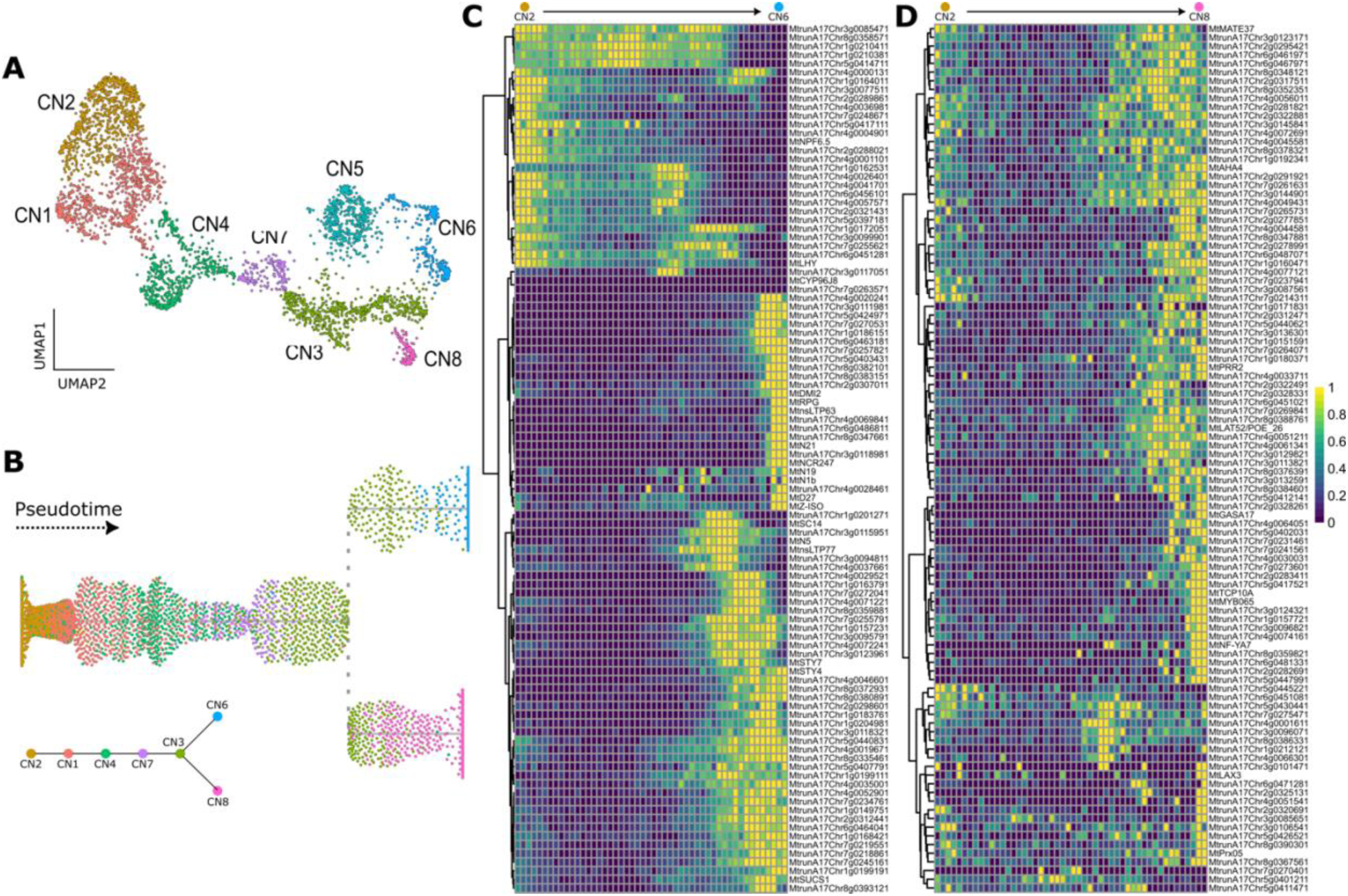
Cell lineages derived from the cortex form the nodule meristem and the infection zone. **(A)** UMAP plot showing the eight groups of cells obtained after reclustering (see Figure 3). **(B)** Cells from those clusters form a bifurcated trajectory containing two lineages. Cells in clusters CN2, CN1, CN4, CN7, CN3, and CN6 are part of the lineage that originates infected cells that will form the ZII of the nodule from the cortical cells. Cells in clusters CN2, CN1, CN4, CN7, CN3, and CN8 are part of the lineage originating the nodule meristem from cortical cells. **(C)** The 100 most significant DEGs (TradeSeq, FDR < 0.001, sorted by the Wald test statistic) within the trajectory from the cortex (CN2) to infected cells (CN6). **(D)** The expression of the most significant DEGs (same parameters as in C) within the trajectory from the cortex (CN2) to meristem (CN8). Only DEGs exclusively detected in one of the two trajectories are shown. The complete list of DEGs, including the DEGs detected in both trajectories, is shown in Additional file 3. Heatmaps were generated as described previously (Figure 2).

Based on the LCM data, we inferred that CN1 and CN2 are composed of cortical cells (Figure S15) – these clusters were detected throughout the experiment (0 to 96 hpi, Figure S12- B). Among these cells, those in CN2 were significantly enriched for the expression of CHASE- domain containing histidine kinase receptors (*MtCHK3* and *MtCHK4*) and other cytokinin signaling genes (*e.g.*, *MtRRB24*). Interestingly, this group of cells was also enriched for several HD-ZIP I transcription factors, including *MtHB1* and other Homeobox-WOX gene family members (*MtrunA17Chr5g0400781*; *MtrunA17Chr3g0123691*). In Medicago lateral root development, *MtHB1* represses a LOB-like (for Lateral Organ Boundaries) gene, *LBD1*, which is auxin-inducible during lateral root emergence (Ariel *et al*., 2010).

Markers for the nodule primordia such as *BASIC HELIX-LOOP-HELIX 1* (*MtbHLH1*, Godiard *et al*., 2011) (Figure 3C) and *AUXIN INFLUX TRANSPORTER LAX2* (*MtLAX2*, Roy *et al*., 2017) were strongly expressed in CN4, and part of CN3 and CN7. CN3 was particularly enriched for auxin biosynthesis, transport, and signaling genes. Among the genes specifically expressed in this cluster are several components of the *SHORT INTERNODES/STYLISH* (STY) family (*MtSTY3*, *MtSTY4,* and *MtSTY7*). LOB-domain proteins regulate these genes and, together with genes of the *YUCCAs* (YUC) family, participate in generating a local auxin maximum required for the initial nodule developmental program (Schiessl *et al*., 2019). The same cluster- specific genes included auxin influx and auxin efflux carriers (*e.g.*, *MtPIN2* and *MtLAX2*). Finally, the expression of members of the *LIGHT SENSITIVE SHORT HYPOCOTYL* transcription factor gene family (*LSH1* and *LSH2*), which have been recently shown to the required for the specification of nodule primordia (Schiessl *et al*., 2023), was also enriched in CN3 (*LSH1* and *LSH2*) and CN7 (*LSH1*).

Markers for nodule meristem, including *MtWOX5* (Franssen *et al*., 2015), the *PLETHORAS MtPLT3* and *MtPLT5* (Franssen *et al*., 2015), and *MtNOOT2* (Magne *et al*., 2018) were expressed in CN3 and CN8 (Figure 3C). Analysis of the expression of the most specific genes of these clusters in two reference datasets (Roux *et al*., 2014; Schiessl *et al*., 2019) reinforces this annotation (Figure 3D). Genes specific to CN3 and CN8 were more highly expressed in infected than in control samples at 48 hpi in a bulk RNA sequencing dataset (Schiessl *et al*., 2019) and were detected primarily in the fraction of the nodule that contains the meristem (FI; Roux *et al*., 2014) (Figure 3D).

In addition to the cells involved in the nodule meristem development, the infected cells originating from the cortical cells were also identified, represented by CN6 (Figure 3). Among the most specifically expressed genes in this cluster (Additional file 2), most encoded for proteins previously associated with RNS, such as nodulins and early nodulins (*MtN1b*, *MtN6*, *MtN7*, *MtN19*, *MtN20*, *MtN21*, *MtN26-4*, *MtENOD12*, and *MtENOD16*), NCRs (*MtNCR057*, *MtNCR247*, *MtNCR514*, *MtNCR657*), NFYs (*MtNFY-A1*, *MtNF-YB16*, *MtNF-YB18*), *NPL*, *NODULE-INDUCED RECEPTOR-LIKE KINASE 1* (*MtNRLK1*), *MtVPY*, *MtRPG*, *MtRRA3*, *SYMBIOTIC PROTEIN KINASE* 1 (*MtSPK*|*MtKIN2*) and *MtSWEET13*, which are primarily expressed in the nodule infection zone (FIId fraction; Roux *et al*., 2014). Notably, many genes remain uncharacterized and whose expression is specific to those infected cells. For many of them, changes in the expression profile during RNS have also been captured in other published datasets, as shown in Figure 3D. Now that the specificity of their expression has been shown at the single-cell level, pointing to a specific role in the infected cortical cells, they may represent candidates for further investigation.

### Trajectory analysis reveals cell lineages that emerge from the cortex to form the nodule compartments

Altogether, annotating these clusters points to a transitory state of cells from CN2 (cortex) to the cells forming the nodule meristem (part of CN3 and CN8). Likewise, a transition from cortical cells (CN2) to the cortical-derivate infected cells (CN6) is also captured. This data suggests that CN3 harbor cells on the verge of differentiating to become the meristem or the infection zone. To evaluate this hypothesis, developmental trajectories were inferred using pseudotime analysis by applying Slingshot (Street *et al*., 2018) (Figure 4), considering as a starting point the cluster CN2. Noteworthy, two connected lineages were uncovered (Figure 4B), with CN3 constituting the branching point separating these lineages. Detection of these lineages agrees with a developmental map of nodule formation in *M. truncatula*, where both meristem and infected cells originate from the root cortex (Xiao *et al*., 2014).

The shared path of the trajectory (CN2 to CN3) shows that it might not be possible to distinguish the dividing cortical cells transcriptionally originated from the separated cortical layers until they reach a more specialized stage during the nodule development. DEGs within the trajectory of each lineage were identified for a total of 5,733 DEGs (Additional file 3). From those, 1,495 were detected in both lineages, 4,010 were exclusively identified in the lineage from cortex to infected cells (CN2 to CN6), and 228 in the lineage from cortex to meristem (CN2 to CN8).

Among the most significant DEGs shared in both lineages are genes well known for their role in root nodule symbiosis, including *SCARECROW* (*MtSCR*), *ANNEXIN 1* (*MtAnn1*), and *MtN18* (Additional file 3). Several more critical genes in the RNS are among the most significant DEG exclusively detected on the lineage from cortex to infected cells (CN2 to CN6), such as *MtRPG*, *DOES NOT MAKE INFECTION 2* (*MtDMI2*), *MtN5*, *MtN19*, *MtN21*, *DWARF27* (*MtD27*) and *MtNCR247*, as well as genes involves in auxin biosynthesis whose function in nodulation has not been verified (*e.g.*, *MtSTY7* and *MtSTY4*) (Figure 4C). A smaller number of exclusive DEGs was identified for the lineage from cortex to meristem, most of which have not been characterized. Among the characterized genes, the transcription factor *MtTCP10A* is one of the most significant (Figure 4D). Its downregulation (with *MtTCP3* and *MtTCP4*, all targets of *MtmiR319*) has been shown to affect the number of nodules in plants overexpressing *MtmiR319* (Wang *et al*., 2018).

Besides the clusters mentioned above, there is a cluster of cells (CN5) for which it was not possible to identify its cell type. While Monocle’s algorithm identified genes as specifically expressed in this cluster, the evaluation of their expression (Figure 3D) shows that most of those genes are, in fact, also expressed in other clusters. Therefore, CN5 was assumed to be a mixture of cell types and was not further explored.

## DISCUSSION

Here we present a single-cell transcriptome dataset encompassing multiple developmental stages of RNS in Medicago. Combined with the inclusion of a hypernodulating mutant, this dataset expands the capture of cells responding to the infection compared to wild-type genotypes. Our data show that gene expression along the cell lineages forming a nodule involves an interplay between the phytohormones auxin and cytokinin, where activation of one is often accompanied by mechanisms to suppress the other (Figure 5). The establishment of a nodule primordium requires the formation of auxin maxima, triggering a developmental program that is similar to that of lateral root formation (Schiessl *et al*., 2019). We show that the specific expression of CK receptors (*MtCHK3*, *MtCHK4*) in the cortex cell cluster CN2 is paralleled by the transcriptional activation of *MtHB1*. As *MtHB1* suppresses the expression of *MtLBD1,* involved in the establishment of an auxin maximum and lateral root formation in Medicago (Ariel *et al*., 2010). Thus, it may play a similar role in limiting the establishment of cell lineages that results in nodule formation. The decrease in expression of *MtHB1* in the next steps of the cell lineage forming a nodule (cluster CN3) was followed by the broad activation of a suite of auxin biosynthesis (*MtSTY3*, *MtSTY4* and *MtSTY7*) and transport genes (*MtPIN2*, *MtLAX2*, *MtABAB29*) supporting the critical function of this phytohormone and its transport in the development of the nodule primordium. Finally, the specific activation of a suite of nodulation genes (*e.g.*, *NIN*, *NF-YA1*, *NF-YB16* & *NF-YB18*) in the developing nodule (CN6) was accompanied by a highly specific expression of several genes involved in the inactivation (*MtAPT1* and *MtAPT2;* Zhang *et al*., 2013) and degradation of cytokinin (*MtCKX3*, *MtCKX7;* Schmülling *et al*., 2003). While the activation of several nodulation genes may require CK in the initial stages of nodule development (van Zeijl *et al*., 2015), the data suggests suppression as the distinct compartments of the nodule become formed.

**Figure 5.**
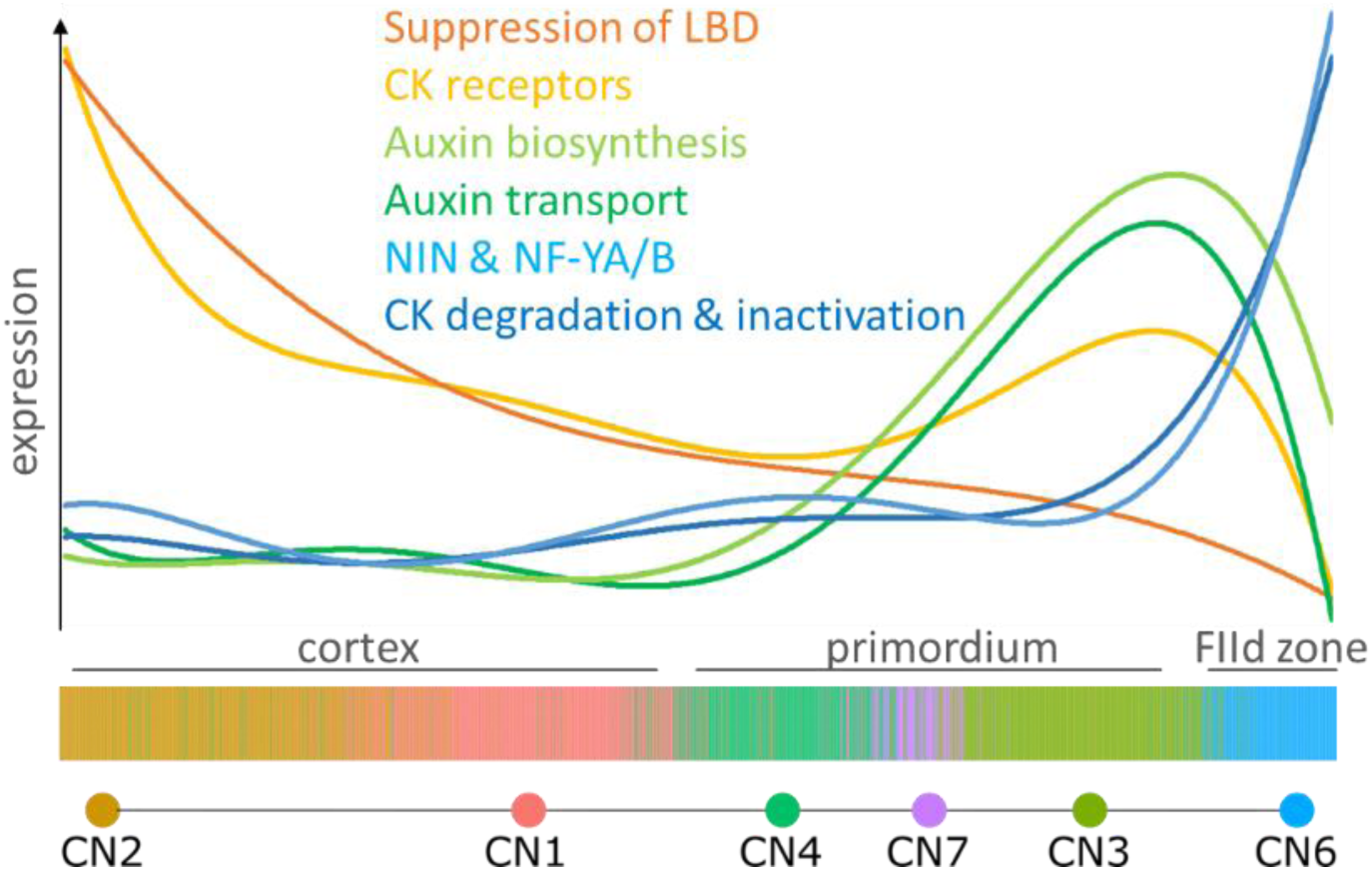
Relative gene expression along the cell lineage trajectory from cortex to the nodule infection (FIId) zone. Gene expression in cells along the trajectory (from CN2 to CN6) were normalized, scaled (0 to 1) and binned as described previously (Fig. 2). To facilitate visualization of gene expression patterns along the trajectory, logarithmic curves were fit based on the average of genes in each category, including suppression of LBD (*MtHB1*); CK receptors (*MtCHK3*, *MtCHK4*), auxin biosynthesis (*MtSTY3*, *MtSTY4*, *MtSTY7*), auxin transport (*MtPIN2*, *MtLAX2*, *MtABAB29*), CK degradation and inactivation (*MtAPT1*, *MtAPT2*, *MtCKX3*, *MtCKX7*) and the nodulation genes *MtNIN*, *MtNF-YA1*, *MtNF-YA2*, *MtNF-YB16*, *MtNF-YB18*.

The high-resolution of single-cell data also allowed us to partition the role of RNS gene family members along cortex-derived cell lineages or epidermal and pericycle cells responding to the rhizobia. The nodule developmental program involves genes that are often part of large gene families for which the role of only a few members has been hypothesized or validated in RNS. The resource described here provides a quantitative assessment of the transcription of individual family members. While the functional validation of the function of genes identified in association with RNS is beyond the scope of this study, it establishes hypothesis about the redundancy or complementary roles specific gene family members at specific cell types. For instance, the LOB domain protein gene *MtLBD16* function in nodulation has been extensively characterized (Schiessl *et al*., 2019) – a role supported by our observation that its expression in cortical cells is amplified after infection, expanding to the nodule primordium (C16) and developing nodule (C6; Figure S16). However, the high expression of the uncharacterized *MtLBD4* and *MtLBD18* during RNS suggests a similar role in this symbiosis (Figure 6). Before infection, *MtLBD4* expression is specific to the stele and pericycle but rapidly (<24 hpi) expands to the infected root hair cells, remaining high until at least 96 hpi. At this latest time, *MtLBD4* is also highly expressed in the nodule (C6) (Figure S16). *MtLBD18* follows an expression pattern similar to *MtLB16* in the cortex and nodule, but its expression is significantly higher in those clusters (Figure S16). The detection of *LBD*s in the infected root hair cells (Figure 6) also suggests a possible role of specific gene family members during root hair infection, in addition to the previously described function in nodule organogenesis.

**Figure 6.**
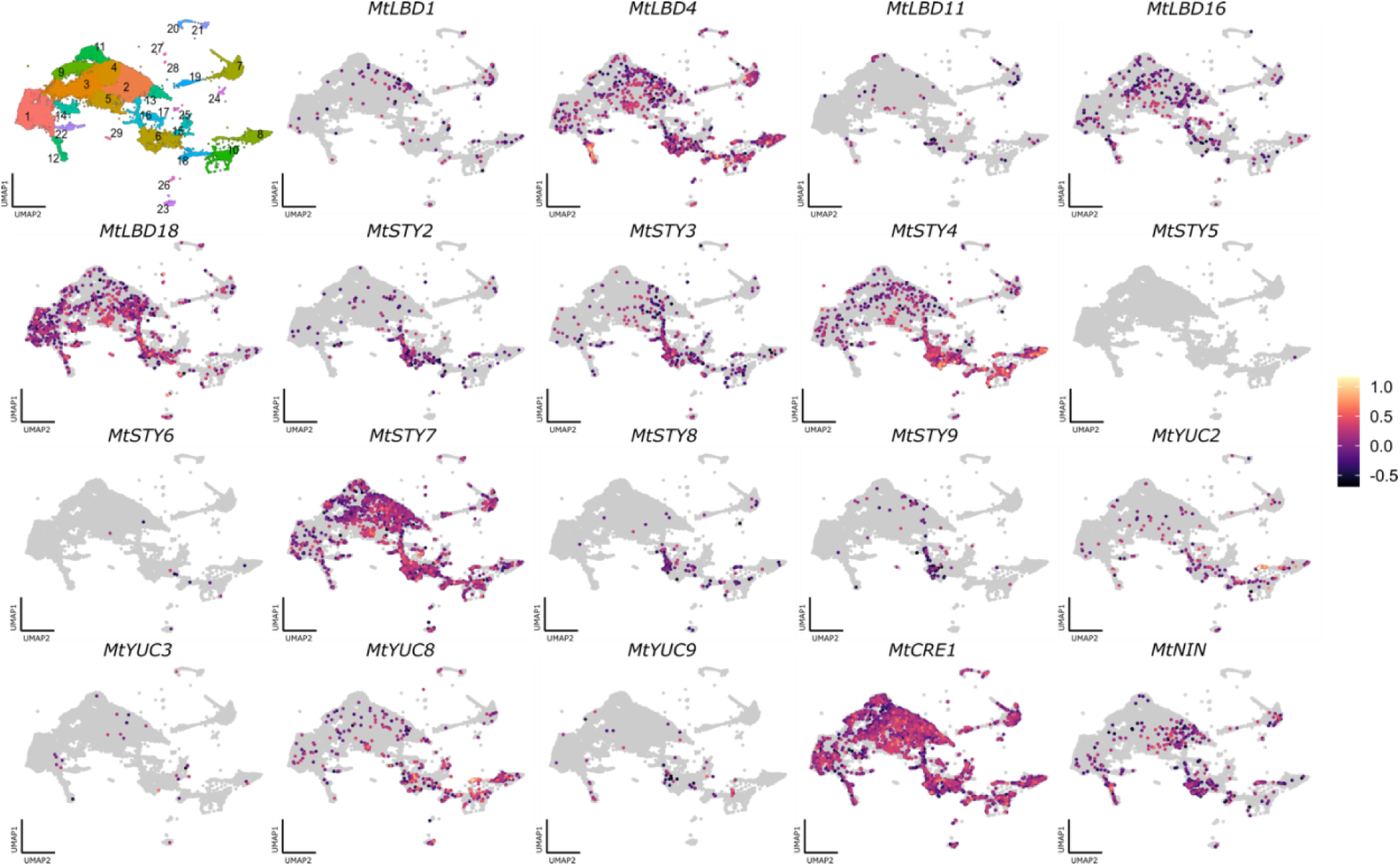
Expression profile of all annotated *LBD* and *STY* genes in the *M. truncatula* genome (v.5, release 1.9). The UMAP plots represents the cells captured in all time points and in both A17 and *sunn-4* genotypes. Each UMAP plot shows the log10 of the expression (default setting in Monocle3). The expression of *MtCRE1* and *MtNIN*, key regulatory elements of RNS, is shown for comparison.

The single-cell data reported in this resource also provides a spatiotemporal representation of genes implicated in auxin biosynthesis, transport, and signaling in RNS. *LBD*s mediate the activation of *STYLISH* and other regulators of auxin biosynthesis (Schiessl *et al*., 2019), of which only a few have been characterized for their role in RNS. We found that most genes of the eight-member STY family were expressed during nodule development, except for *MtSTY5* and *MtSTY6*. While there was a high degree of spatial and temporal expression similarity among some members (for example, *MtSTY2* and *MtSTY3*, and *MtSTY8* and *MtSTY9*), unique patterns of transcript abundance also suggest complementary roles in RNS. For instance, except for *MtSTY4* and *MtSTY7*, none of the STY were expressed before rhizobial infection in the root (Figure S16). Under control conditions (0 hpi), *MtSTY4* was expressed primarily in the stele (C10) and the pericycle (C8), while *MtSTY7* expression was more abundant in cortical cells throughout the experiment. These and all other STYs expressed during RNS were detected in clusters related to nodule development (C16 and C6) and in the small subset of cells from pericycle that appear after the infection (C8) at 96 hpi (Figure 6 and S16).

Finally, the analysis reported here identified many genes whose expression is activated following rhizobial infection, in lineages that emerge during nodule development or in the infected cells of the root hair and infection zone. While further investigation and experimental validation will be necessary to verify those findings, accessible exploration of the expression profile of all genes investigated in our dataset (https://) will accelerate their characterization.

## Acknowledgments

We would like to thank Thomas B. Irving, currently at University of Cambridge, for providing a list of genes involved in the root nodule symbiosis.

## Authors contribution

Conceptualization, CD, DC, FAF, JF, JMA, KMB, MK, PMT, SR and WJP; Methodology, MK and WJP; Software, WJP; Validation, WJP; Formal Analysis, JB, KMB and WJP; Investigation, CN, CD, DC, HWS, JB, PMT and YG; Resources, JF, JMA, MK and SR; Data Curation, CD, MK and WJP; Writing – Original Draft, CD, JB, MK and WJP; Writing – Review & Editing, CN, CD, DC, FAF, HWS, JB, JF, KMB, MT, MK, NCP, PMT, SC, SAK, SR, TBI, YG and WJP; Visualization, JB, DC, MK, YG and WJP; Supervision, CD, JF, JMA, MK and SR; Project Administration, CD, MK, and WJP; Funding Acquisition, JF, FAF, JMA, MK and SR.

## DECLARATIONS

### Ethics approval and consent to participate

Not applicable.

### Consent for publication

Not applicable.

## Data and code availability

### Funding

We thank the United States Department of Energy, Office of Science, Biological and Environmental Research program, for funding this study (DE-SC0018247 to MK, SR, and J-MA), and Clemson University SUCCEEDS to JF and FAF.

## METHODS

### Plant material and growth conditions

In this study, two genotypes of *M. truncatula* were used, Jemalong A17 and a mutant for the gene *SUNN*, namely *sunn-4* (Schnabel *et al*., 2005). The mutant *sunn-4* presents a hypernodulation phenotype.

*M. truncatula* seeds were scarified and stratified to initiate germination. Scarification was done by soaking seeds in sulfuric acid for 8 min before washing them with water six times. Seeds were sterilized in bleach (12% sodium hypochlorite) for 4 min, washed with sterile water four times, and incubated for 1 h in water. Seeds were sown on 1% agar plates with 1 μM of GA3. The plates were kept at 4 °C for three days to induce stratification. After incubating plates at 24 °C for 12 h in the dark, four seedlings were transferred to CYG germination pouches (Anon, 2014). Each pouch contained four seedlings and 10 mL of Modified Nodulation Medium (Chakraborty *et al*., 2021). Pouches were placed vertically in polypap stands (Anon, 2014), and the stands were placed in a tray and 7.5” (L) x 11’’ (W) x 21.25’’ (H) humidity dome (Alberta LTD, Amazon Standard Identification Number: B089Q24D3N). The inside of the dome was sprayed daily with water to maintain humidity. After 3 days, pouches were replenished with 10 mL of Modified Nodulation Medium. The growth chamber was kept at 24 °C with long-day light conditions (16 h light/8 h dark; 150 μmol m−2 s−1 light intensity).

### Inoculation assay

To prepare the rhizobia for inoculation, a 5 mL liquid culture was grown overnight from a fresh plate of *Sinorhizobium meliloti* 1021. After 100-200 μL of the liquid culture was plated on Tryptone Yeast plates, they were grown for 48 h at 28°C. The entire film of rhizobia was collected in a 50 mL falcon tube and resuspended with autoclaved water. The rhizobia suspension was then diluted to a final OD600 = 0.1 using Fahräeus medium without nitrogen (Boisson-Dernier *et al*., 2001). Plants were grown for one week in pouches before inoculation. Plants were inoculated by pipetting 1 mL of the *S. meliloti* 1021 suspension into each pouch. At the time of inoculation, the susceptibility zone of each plant was marked outside the pouch in the proximity of the root tips.

### Sample Collection

Ten growth pouches were used at each time point for a total of 40 plants each. At each time point, 4 cm sections around the marked susceptibility zone were harvested from each plant. The root segments from the 0 h time point (i.e., before inoculation) served as a control compared with expression data from the later time points. Root segments were harvested at 24, 48, and 96 hpi for each respective time point. The root segments collected at each time point were pooled for nuclei isolation and snRNA-seq.

### Nuclei isolation from *Medicago* roots for single nuclei RNA-seq

The protocol used for nuclei isolation was previously described in (Conde *et al*., 2021) with minor adjustments. Briefly, nuclei isolation was carried out at 4 ℃, and all materials, tools, and solutions were pre-cooled to that temperature. Root segments were moved to a glass plate with 200 μL of modified Nuclei Isolation Buffer (NIB; Conde *et al*., 2021) containing 0.5 U/mL Protector RNase Inhibitor (Sigma Aldrich). After samples were fragmented with a sterile razor blade, the homogenate was washed into a 50 mL tube with NIB and incubated on a shaker for 5 min. All samples underwent at least two filtration stages, first with Miracloth (Calbiochem) and then with a 40 μm cell strainer (Greiner Bio-One). The samples were centrifuged for 5 min at 600 g before the supernatant was discarded without disturbing the pellet. The pellet was resuspended with 4 mL of NIB WASH (Conde *et al*., 2021). Centrifugation and resuspension were repeated once more. After the final centrifugation, 300 μL of NIB WASH was used to resuspend the pellet. Samples that contained debris underwent an additional filtration step with a 40 μm cell strainer with an attached 5 mL tube (Falcon Corning). Next, the suspension was transferred to a sterile tube compatible with the BD FACSAria™ IIU/III upgraded cell sorter. Nuclei were stained with 5 μg/mL DAPI for 5 min at 28 °C before sorting. The sample was used for Florescence Activated Nuclei Sorting (FANS) at the Interdisciplinary Center for Biotechnology Research (ICBR) at the University of Florida to reduce the debris in the sample and minimize the likelihood of clogging the 10× Genomics microfluid chip. Settings for FANS have been described previously (Conde *et al*., 2021).

### Single nuclei cDNA and Library Preparation

To generate snRNA-seq libraries for each genotype and time point, 10× Genomics microfluid chips were loaded with 20 thousand nuclei. The Single Cell v3.1 Dual Index Gene Expression protocol was followed with a few minor adjustments, as outlined in (Conde *et al*., 2021). Briefly, NIB WASH was used to reach the final desired volume of nuclei suspension, and a total of 15 PCR cycles were used to amplify the cDNA. The amplified cDNA was used to construct libraries according to the manfacturer instructions and the cDNA was sequenced at the ICBR using the NovaSeq 6000 System, S1 flow cell 2x100 sequencing kit but with cycling of 28 read 1, 10 index 1, 10 index 2, and 90 read 2.

### Cell clustering

The sequencing output was demultiplexed and processed using the software Cell Ranger (v7.0, 10× Genomics). Cell ranger was also applied to generate counts for each gene in the *M. truncatula* genome (v.5.0, release 1.9; Pecrix *et al*., 2018). Cell ranger was executed using default parameters, except by the inclusion of “--force-cells=10000”, which set the number of cells to be considered by the software to ten thousand.

Combining the samples and clustering was performed using Monocle3, as in commit 87f6e88 at (Anon, 2023). Only cells containing 400 or more detected unique molecular identifiers (UMIs) were selected and used for the analysis. Before clustering, the dataset was corrected for batch effect using monocles’ implementation of the matching mutual nearest neighbors method (Haghverdi *et al*., 2018). Unless mentioned otherwise, the default resolution of 0.0001 was applied during the clustering using the Leiden community detection method (Traag *et al*., 2019).

### Identification of novel *Medicago* cell-type-specific markers

To identify the cell types that constitute each cluster of cells in our sample, we evaluated the expression of cell-type-specific marker genes previously described in the literature. In addition, we identified genes specifically expressed in each cluster and investigated their function as a proxy to define the cell type contained in the cluster. For each cluster, only genes expressed in more than 20% of the cells of the cluster and with an adjusted p-value (q-value) ≤ 0.05, generated by Moran’s I test of spatial autocorrelation, were selected. Moran’s I test measures the dependence of a gene expression on the spatial location defined by the cluster and is effective in finding genes that vary in single-cell RNA-seq datasets (Cao *et al*., 2019). Therefore, it can be interpreted similarly to a differential expression test, identifying genes that vary between groups of cells in the UMAP space. When more than 100 marker genes were detected for a cluster, we selected the 100 genes with higher values for the specificity parameter as markers for that cluster for further exploration (Additional file 2).

Next, we evaluated the expression of those marker genes in a dataset originating from a laser capture microdissection (LCM) experiment investigating the initial hours (0 hpi, 24 hpi, 48 hpi, and 72 hpi) of *M. truncatula* response after infection with *S. medicae* ABS7. Specific expression of a marker gene in a cell type captured by LCM indicates that the cluster represents cells of that type. A similar strategy was deployed to investigate the expression of the top 100 marker genes of each cluster in other relevant datasets publicly available as part of the *M. truncatula* RNA-seq Gene Expression Atlas Project v.2 (Carrere *et al*., 2021).

### Pseudotime analysis

During the response to rhizobia infection, multiple cell types undergo transcriptional regulation promoting developmental changes, for example, in the transition from the different layers of cortical cells to the different zones of the nodule. Pseudotime analysis, can reconstruct the different developmental stages by positioning the cells within an inferred developmental trajectory. Once the trajectory is inferred, it is possible to search for genes whose expression changes along the trajectory, pointing to the potential regulators and other critical genes governing the transition. In this study, we used the Bioconductor packages Slingshot (Street *et al*., 2018) to model the trajectories, and TradeSeq (Van den Berge *et al*., 2020) to identify genes whose expression changes along the trajectory. In addition, we used the toolkit dynverse (Saelens *et al*., 2019) for the visual representation of the trajectories and to identify genes whose expression predicts the inferred trajectory.

