## Supplemental Figures for "The Single-Cell Transcriptome Program of Nodule Development Cellular Lineages in *Medicago truncatula*"

**Running title:** Single-cell analysis of nodule development in Medicago

Wendell J. Pereira[1], Jade Boyd[1], Daniel Conde[1,2], Paolo M. Triozzi[1,2], Kelly M. Balmant[1,3], Christopher Dervinis[1], Henry W. Schmidt[1], Carolina Boaventura-Novaes[1], Sanhita Chakraborty[4], Sara A. Knaack [5], Yueyao Gao () [6,7], Frank Alexander Feltus[6,8,9], Sushmita Roy[5,10,11,12], Jean-Michel Ané[4,12], Julia Frugoli[6,12] and Matias Kirst[1, 12,*]

[1] School of Forest, Fisheries and Geomatics Sciences, University of Florida, Gainesville, USA

[2] Centro de Biotecnología y Genómica de Plantas, Universidad Politécnica de Madrid – Instituto Nacional de Investigación y Tecnología Agraria y Alimentaria, Madrid, Spain

[3] Horticultural Sciences Department, University of Florida, USA

[4] Department of Bacteriology, University of Wisconsin – Madison, Madison, USA

[5] Wisconsin Institute for Discovery, University of Wisconsin, Madison, USA

[6] Department of Genetics & Biochemistry, Clemson University, Clemson, USA

[7] Broad Institute of MIT and Harvard, Cambridge, USA

[8] Biomedical Data Science and Informatics Program, Clemson University, Clemson, USA.

[9] Clemson Center for Human Genetics, Clemson University, Greenwood, USA.

[10] Department of Biostatistics and Medical Informatics, University of Wisconsin, Madison, USA.

[11] Department of Computer Sciences, University of Wisconsin, Madison, USA.

[12] Senior author.

SUPPLEMENTAL FIGURES

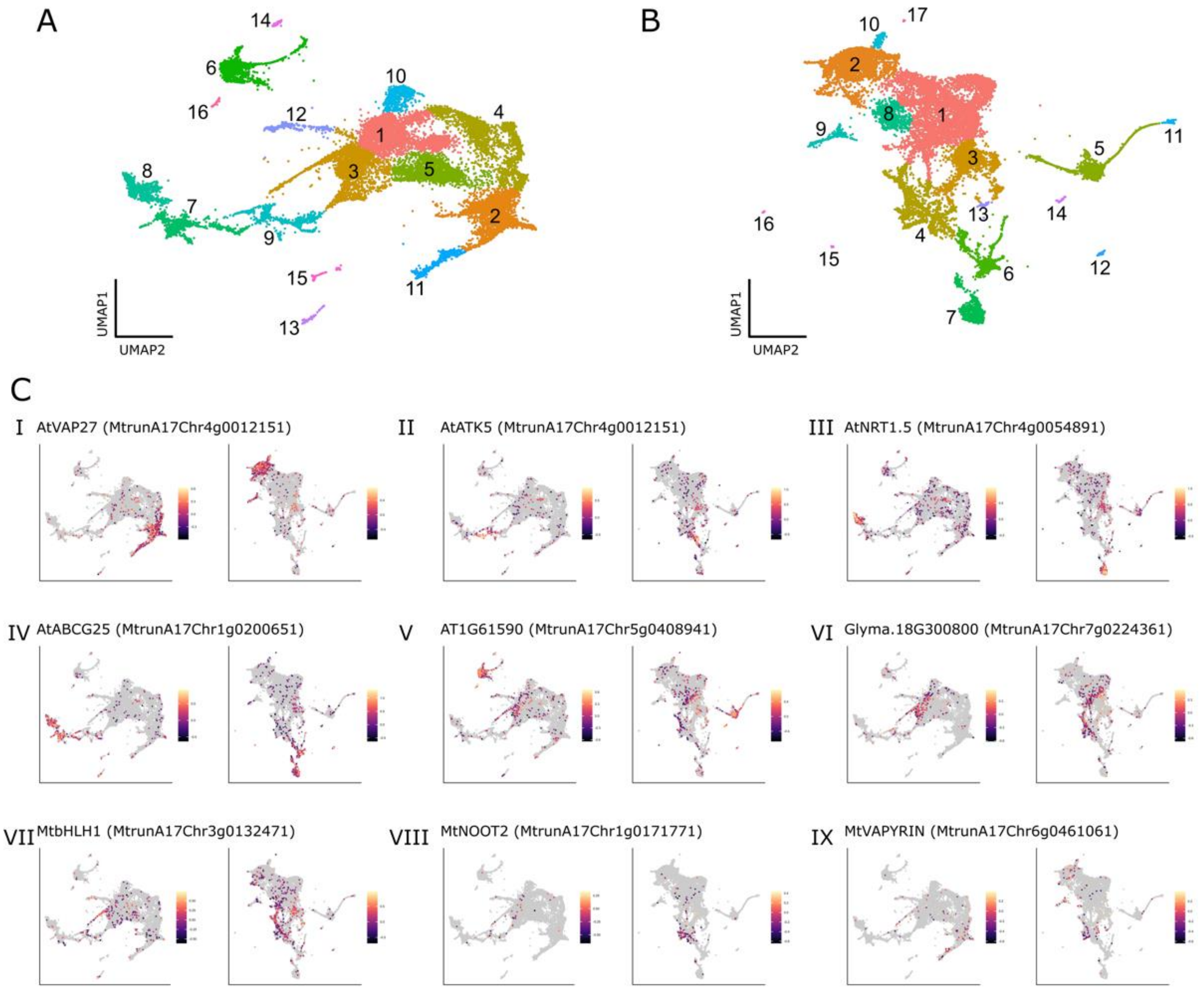


**Fig. S1. Jemalong A17 and *sunn-4* have similar distribution of cell types.** Combining all time points, 16 clusters were obtained for Jemalong A17 (A) and 17 clusters for the *sunn-4* (B) datasets. The expression of marker genes of different cell types confirms their presence in both datasets (C). Selected markers are shown for the epidermis/root hair (I), cell division in the vasculature (II), pericycle (III), stele (IV), endodermis (V), cortex (VI), nodule primordium (VII), nodule meristem (VIII), infection thread (IX). In each UMAP plot, the heatmap in the legend shows the log10 of the expression, which is the default in Monocle3.

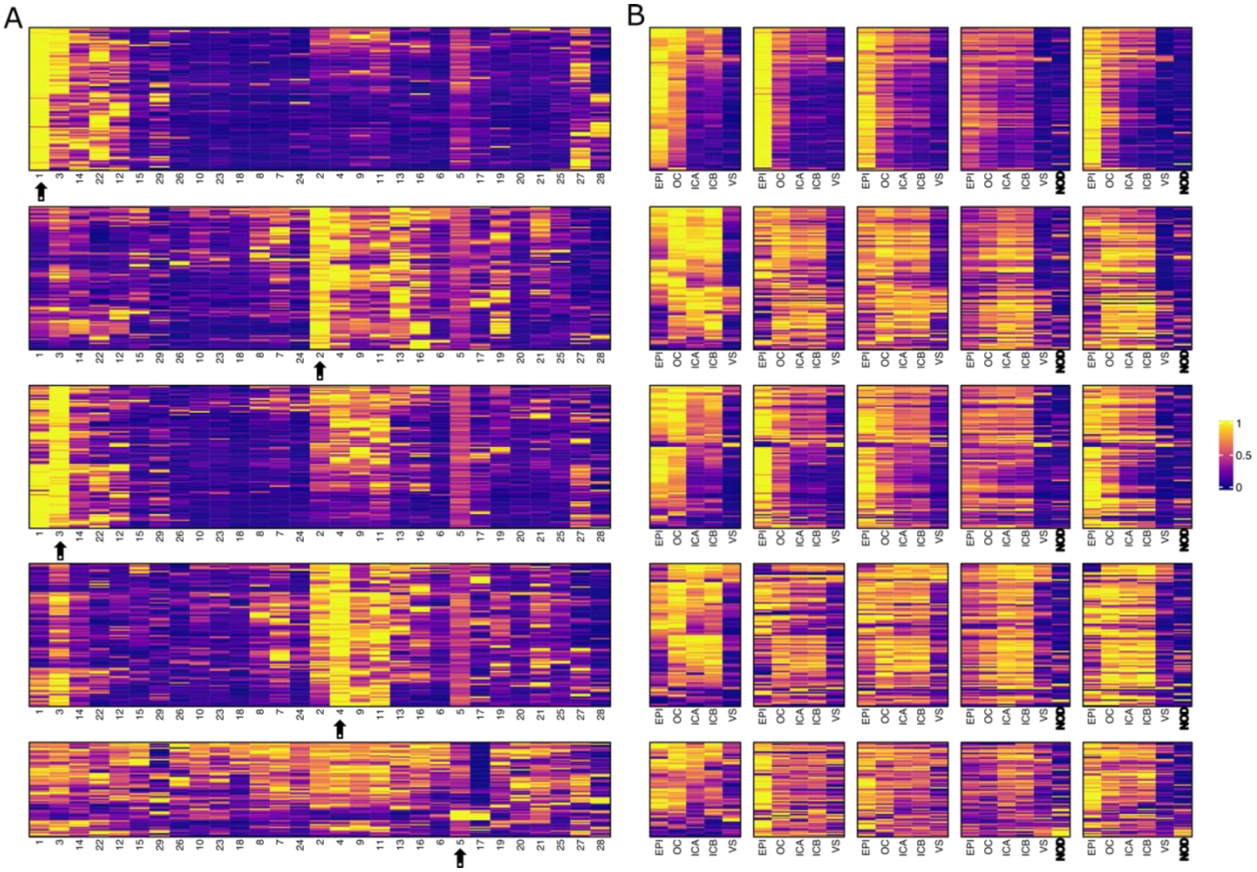


**Fig. S2. Expression profile of the 100 most specifically expressed genes of clusters 1 to 5 and comparison with their expression in the LCM dataset.** In A, the expression profile of the 100 most specific genes (q-value < 0.05, sorted by the specificity value returned by the Moran's I test) of each cluster is shown in comparison with the other clusters in the data. The arrows show which cluster is being represented. Each column shows the average expression of all cells in the cluster. The expression was scaled by row. The clusters within each heatmap follow the order in Figure 1C to reinforce the similar expression among clusters of cells from the same cell type. In B, it is represented the expression profile of the same genes depicted in A in an LCM dataset covering a similar experimental design (see methods). Each column represents the average of three experimental replicates. The expression was scaled by row. EPI = Epidermal cells, OC = Outer cortical cells, ICA = Inner cortical cells at the xylem poles, ICB = Inner cortical cells between the xylem poles, VS = vasculature, NOD = Nodule. Note that nodule cells were only sampled at 48 h and 72 h after infection.

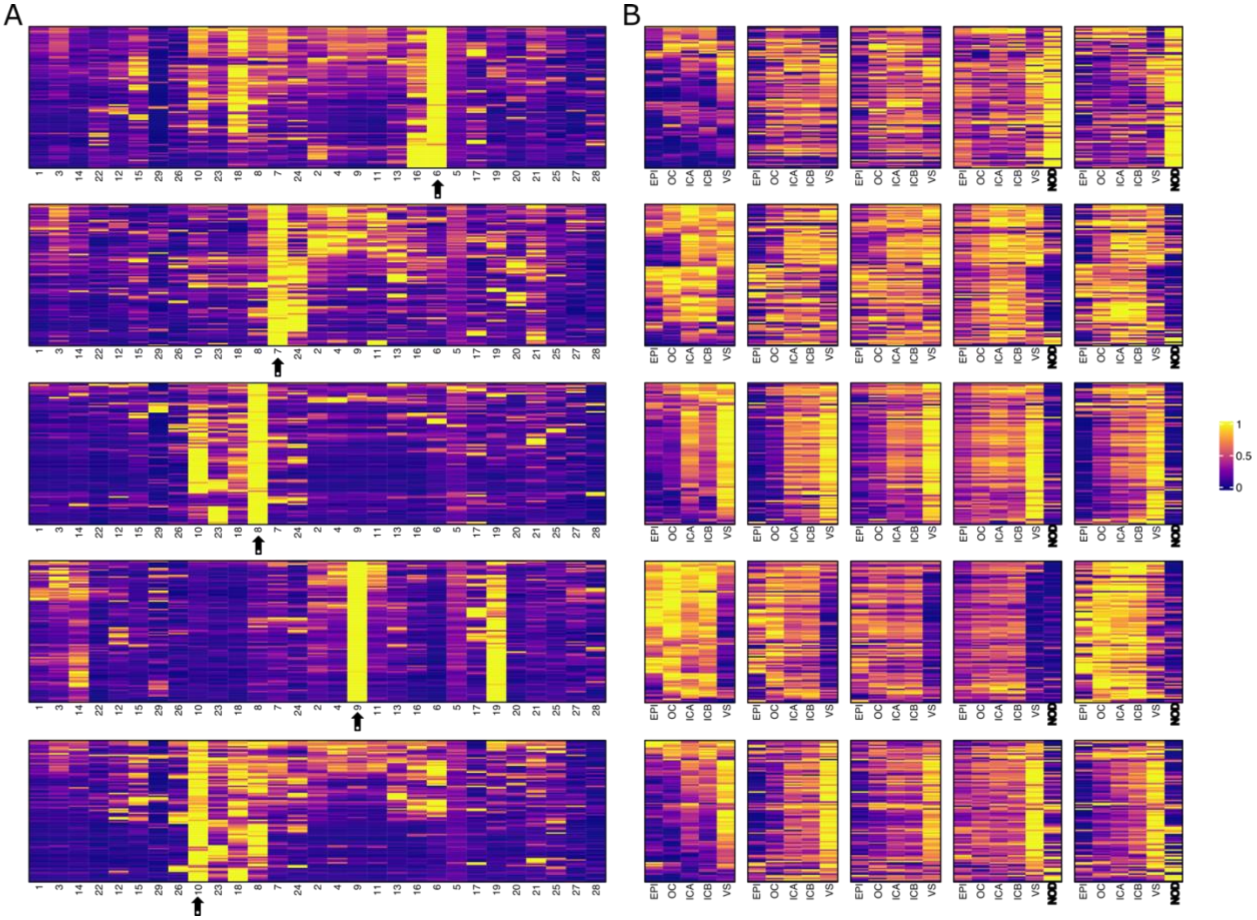


**Fig. S3. Expression profile of the 100 most specifically expressed genes of clusters 6 to 10 and comparison with their expression in the LCM dataset.** In A, the expression profile of the 100 most specific genes (q-value < 0.05, sorted by the specificity value returned by the Moran's I test) of each cluster is shown in comparison with the other clusters in the data. The arrows show which cluster is being represented. Each column shows the average expression of all cells in the cluster. The expression was scaled by row. The clusters within each heatmap follow the order in Figure 1C to reinforce the similar expression among clusters of cells from the same cell type. In B, it is represented the expression profile of the same genes depicted in A in an LCM dataset covering a similar experimental design (see methods). Each column represents the average of three experimental replicates. The expression was scaled by row. EPI = Epidermal cells, OC = Outer cortical cells, ICA = Inner cortical cells at the xylem poles, ICB = Inner cortical cells between the xylem poles, VS = vasculature, NOD = Nodule. Note that nodule cells were only sampled at 48 h and 72 h after infection.

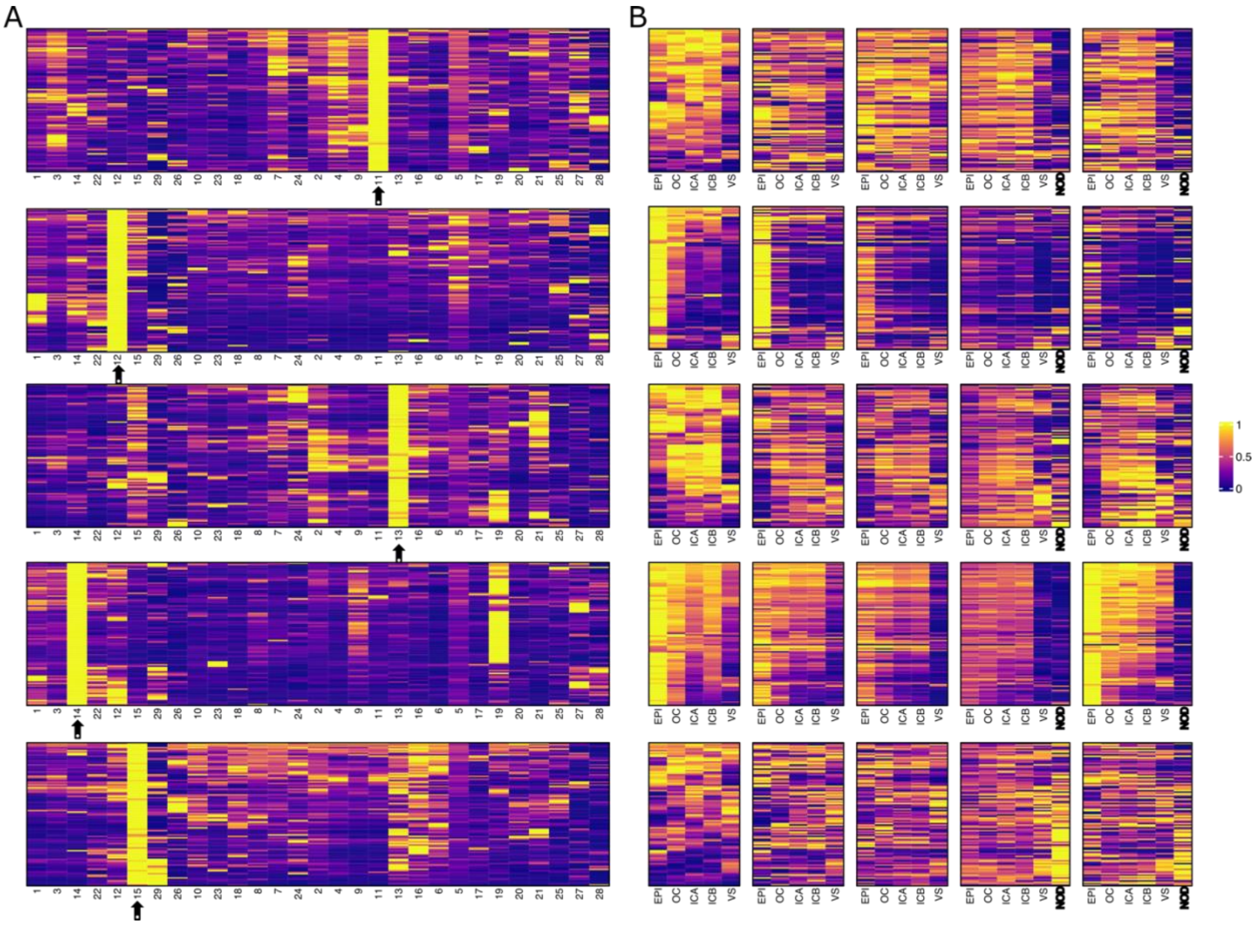


**Fig. S4. Expression profile of the 100 most specifically expressed genes of clusters 11 to 15 and comparison with their expression in the LCM dataset.** In A, the expression profile of the 100 most specific genes (q-value < 0.05, sorted by the specificity value returned by the Moran's I test) of each cluster is shown in comparison with the other clusters in the data. The arrows show which cluster is being represented. Each column shows the average expression of all cells in the cluster. The expression was scaled by row. The clusters within each heatmap follow the order in Figure 1C to reinforce the similar expression among clusters of cells from the same cell type. In B, it is represented the expression profile of the same genes depicted in A in an LCM dataset covering a similar experimental design (see methods). Each column represents the average of three experimental replicates. The expression was scaled by row. EPI = Epidermal cells, OC = Outer cortical cells, ICA = Inner cortical cells at the xylem poles, ICB = Inner cortical cells between the xylem poles, VS = vasculature, NOD = Nodule. Note that nodule cells were only sampled at 48 h and 72 h after infection.

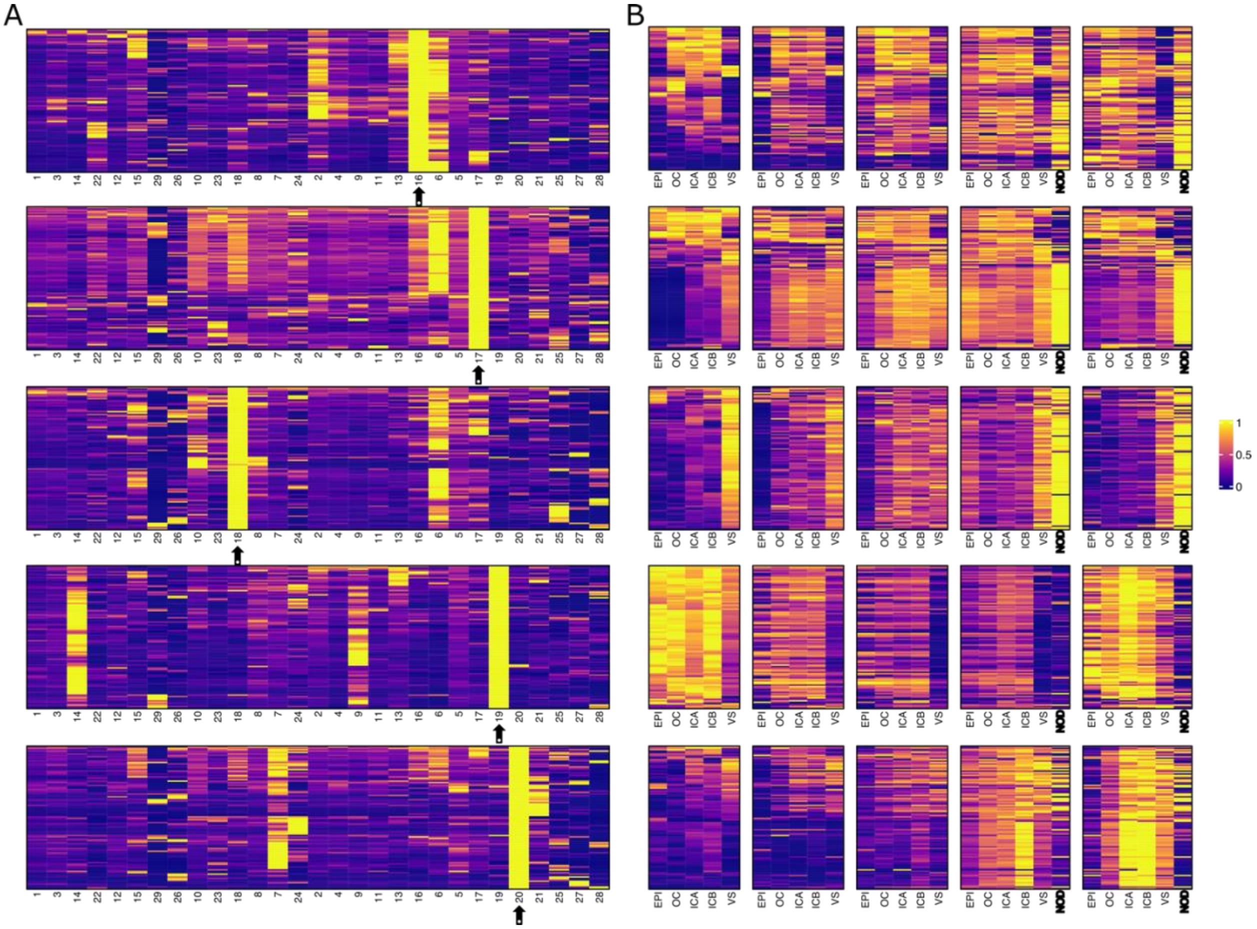


**Fig. S5. Expression profile of the 100 most specifically expressed genes of clusters 16 to 20 and comparison with their expression in the LCM dataset.** In A, the expression profile of the 100 most specific genes (q-value < 0.05, sorted by the specificity value returned by the Moran's I test) of each cluster is shown in comparison with the other clusters in the data. The arrows show which cluster is being represented. Each column shows the average expression of all cells in the cluster. The expression was scaled by row. The clusters within each heatmap follow the order in Figure 1C to reinforce the similar expression among clusters of cells from the same cell type. In B, it is represented the expression profile of the same genes depicted in A in an LCM dataset covering a similar experimental design (see methods). Each column represents the average of three experimental replicates. The expression was scaled by row. EPI = Epidermal cells, OC = Outer cortical cells, ICA = Inner cortical cells at the xylem poles, ICB = Inner cortical cells between the xylem poles, VS = vasculature, NOD = Nodule. Note that nodule cells were only sampled at 48 h and 72 h after infection.

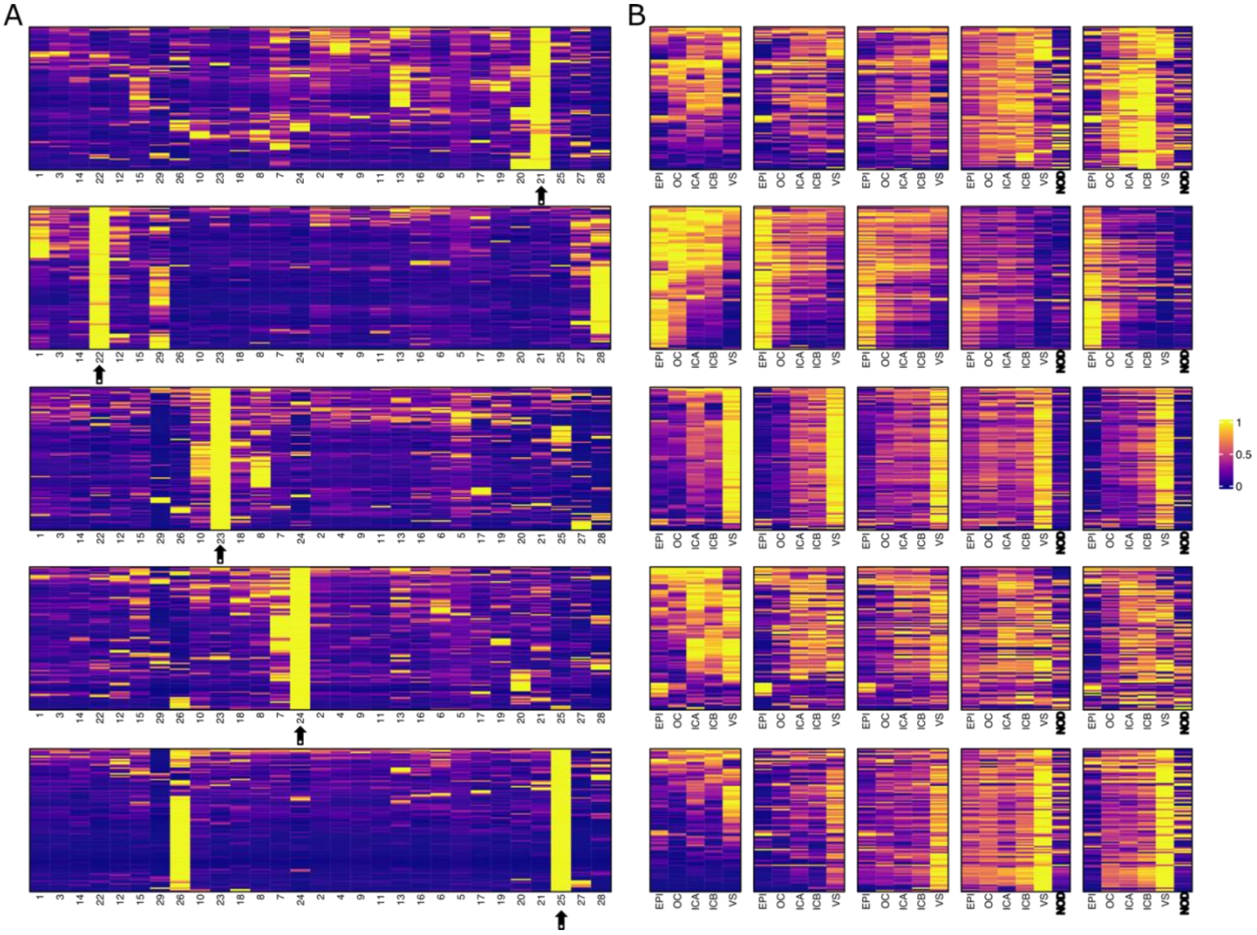


**Fig. S6. Expression profile of the 100 most specifically expressed genes of clusters 21 to 25 and comparison with their expression in the LCM dataset.** In A, the expression profile of the 100 most specific genes (q-value < 0.05, sorted by the specificity value returned by the Moran's I test) of each cluster is shown in comparison with the other clusters in the data. The arrows show which cluster is being represented. Each column shows the average expression of all cells in the cluster. The expression was scaled by row. The clusters within each heatmap follow the order in Figure 1C to reinforce the similar expression among clusters of cells from the same cell type. In B, it is represented the expression profile of the same genes depicted in A in an LCM dataset covering a similar experimental design (see methods). Each column represents the average of three experimental replicates. The expression was scaled by row. EPI = Epidermal cells, OC = Outer cortical cells, ICA = Inner cortical cells at the xylem poles, ICB = Inner cortical cells between the xylem poles, VS = vasculature, NOD = Nodule. Note that nodule cells were only sampled at 48 h and 72 h after infection.

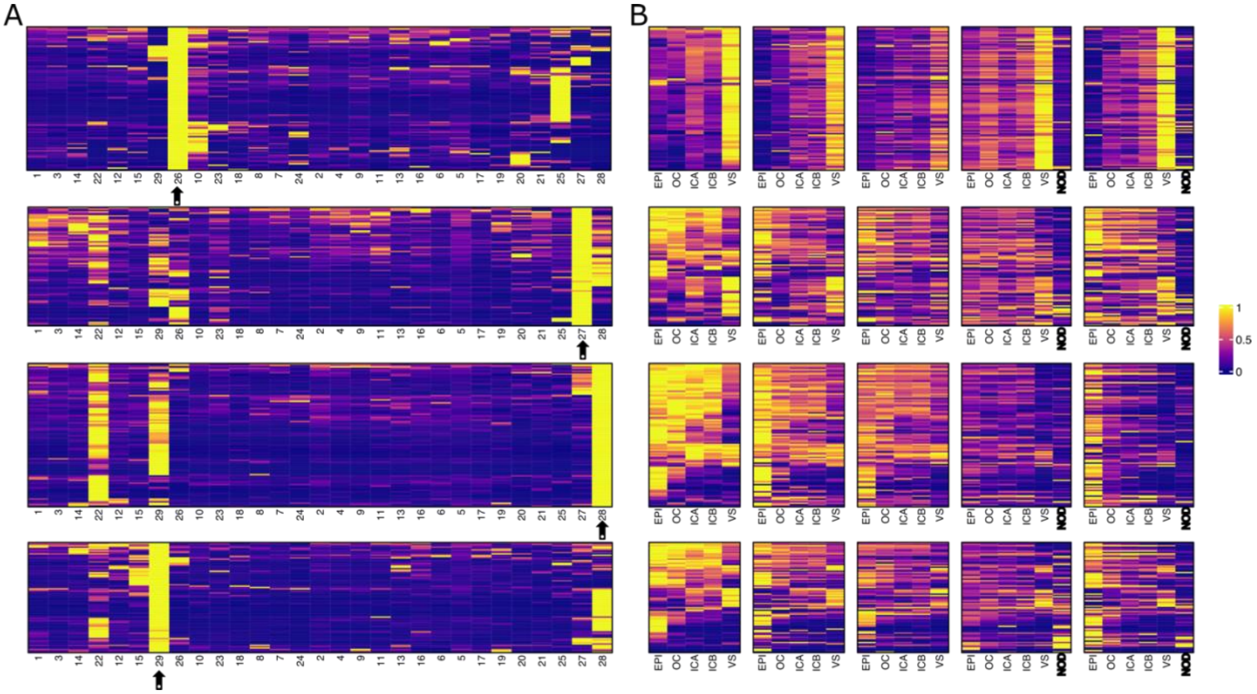


**Fig. S7. Expression profile of the 100 most specifically expressed genes of clusters 25 to 29 and comparison with their expression in the LCM dataset.** In A, the expression profile of the 100 most specific genes (q-value < 0.05, sorted by the specificity value returned by the Moran's I test) of each cluster is shown in comparison with the other clusters in the data. The arrows show which cluster is being represented. Each column shows the average expression of all cells in the cluster. The expression was scaled by row. The clusters within each heatmap follow the order in Figure 1C to reinforce the similar expression among clusters of cells from the same cell type. In B, it is represented the expression profile of the same genes depicted in A in an LCM dataset covering a similar experimental design (see methods). Each column represents the average of three experimental replicates. The expression was scaled by row. EPI = Epidermal cells, OC = Outer cortical cells, ICA = Inner cortical cells at the xylem poles, ICB = Inner cortical cells between the xylem poles, VS = vasculature, NOD = Nodule. Note that nodule cells were only sampled at 48 h and 72 h after infection.

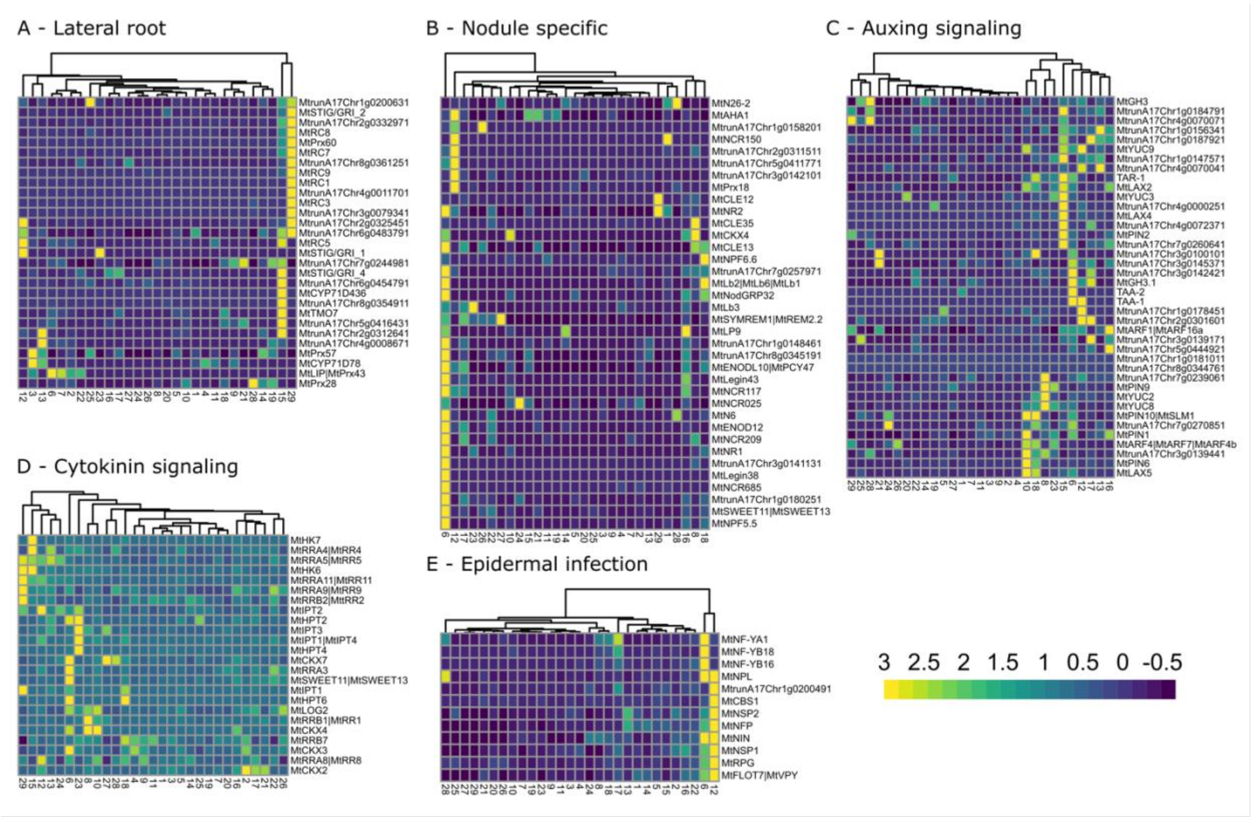


**Fig. S8. Expression profile of genes reported by Schiessl et al., 2019.** The shown genes were selected due to their specific expression in the lateral root (A) and nodules (B), or due to their role during the auxin (C) and cytokinin (D) signaling or epidermal infection (E).

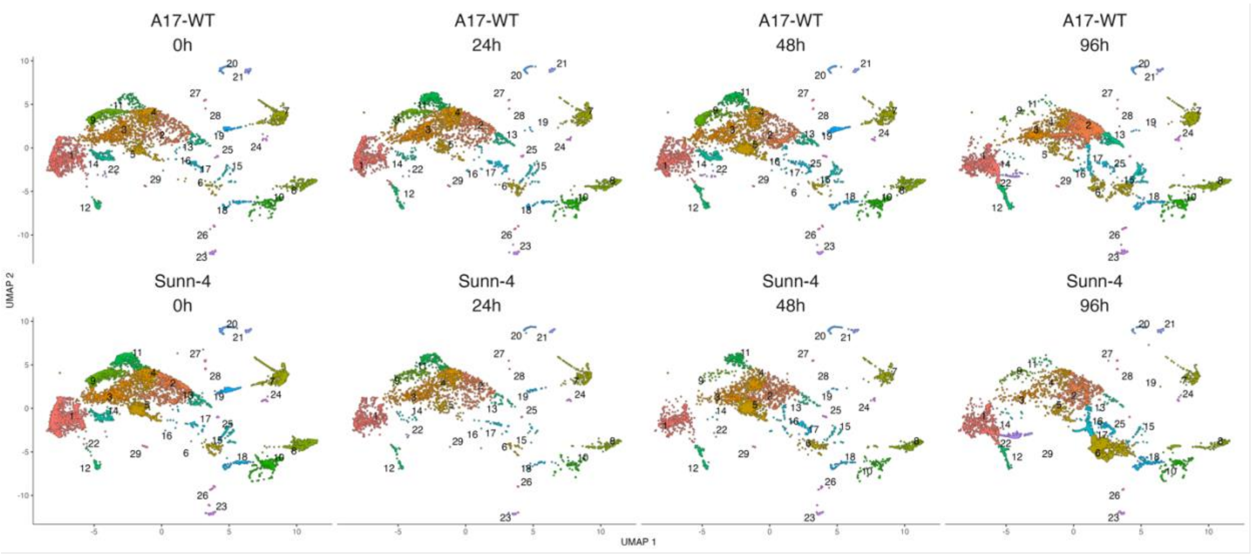


**Fig. S9. Changes in the distribution of cells among clusters in response to the rhizobia infection.** From left to right, the plots show the distribution of cells at 0h, 24h, 48h, and 96 after the infection. On the top are the UMAP plots of Jemalong A17, and on the bottom are the UMAP plots of *sunn-4*.

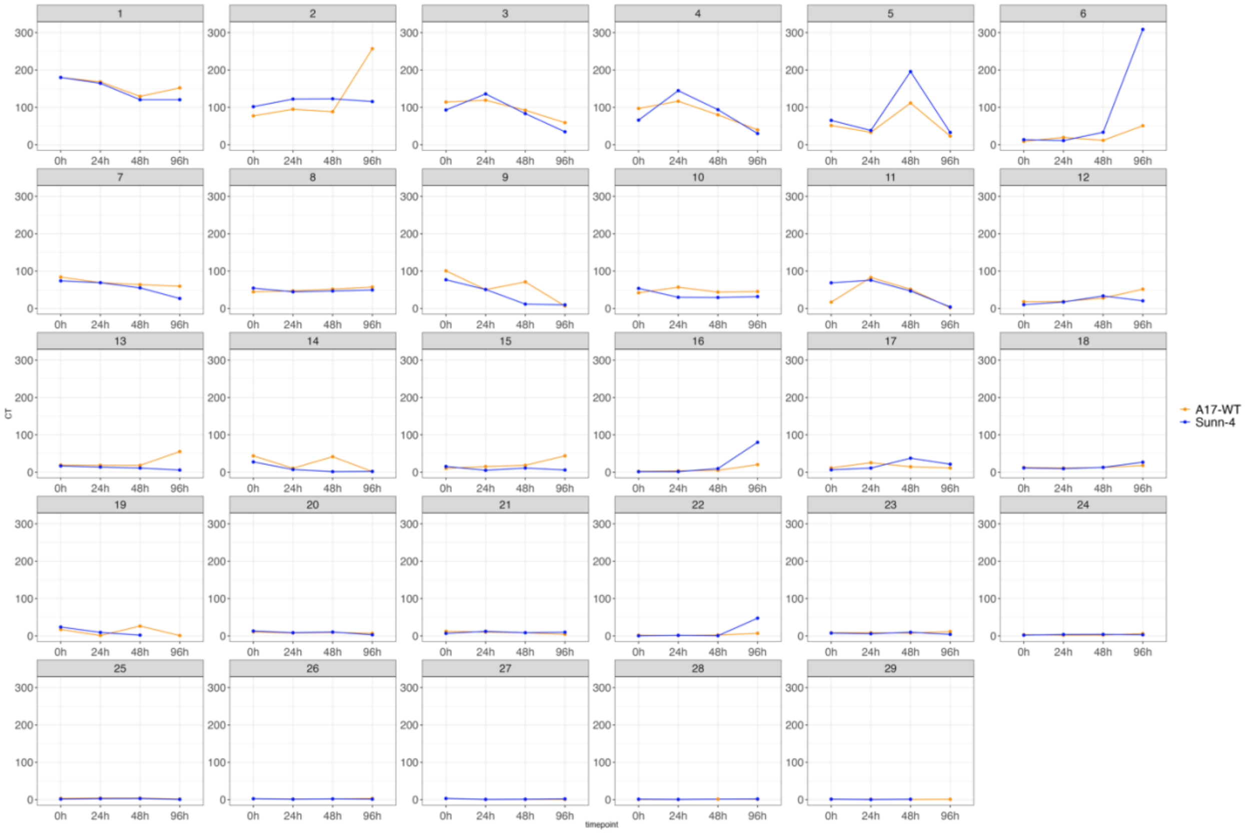


**Fig. S10. Number of cells per cluster and genotype, according to the time after infection.** To avoid bias due to the number of sampled cells in each sample, the y-axis represents the number of cells per thousand cells in each sample.

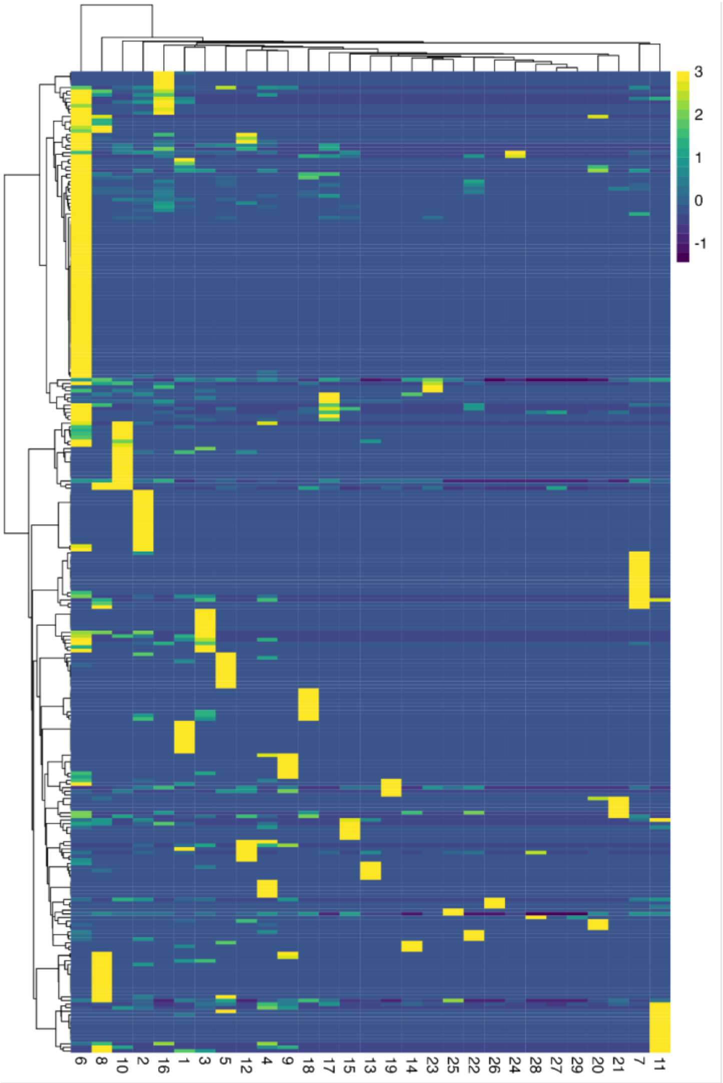


**Fig. S11. Expression profile of the 272 Nodule Cysteine-Rich peptides detected in this dataset.** Genes (rows) and clusters (columns) were grouped by a hierarchical clustering algorithm. The names and averaged expressions of each NCR per cluster are available in Table S1.

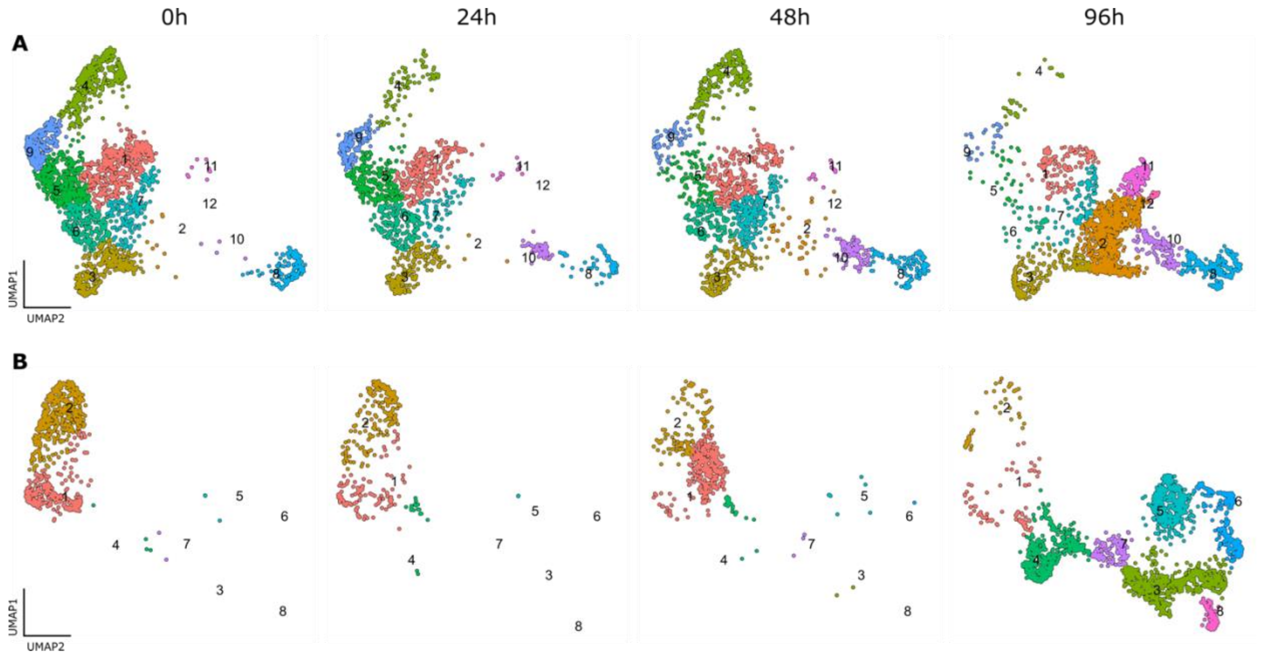


**Fig. S12. Changes in the distribution of cells among clusters in response to the rhizobia infection after the reclustering of specific cell types.** Distribution of cells after reclustering cells from root hair and epidermis (A) and cells from cortex and nodule (B). From left to right, the plots show the distribution of cells at 0h, 24h, 48h, and 96h after the infection. In A, the plots show cells from Jemalong A17 and *sunn-4*. In B, only cells from the *sunn-4* genotype were used.

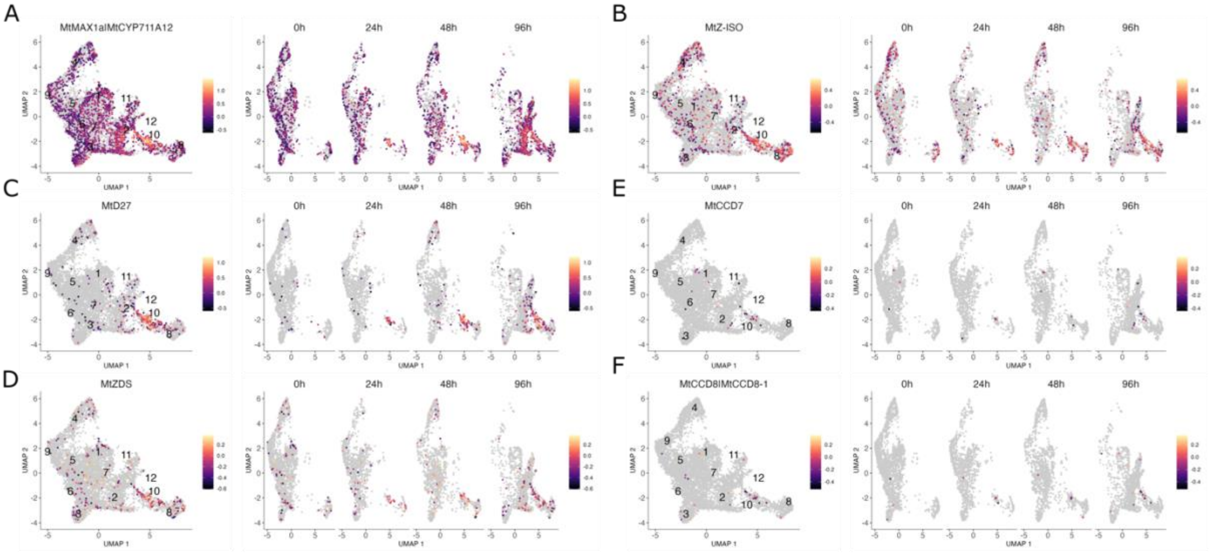


**Fig. S13. Expression profile of genes in the Strigolactones biosynthesis pathway in the root hair reclustered dataset.** In each UMAP plot, it is shown the log10 of the expression, which is the default in Monocle3.

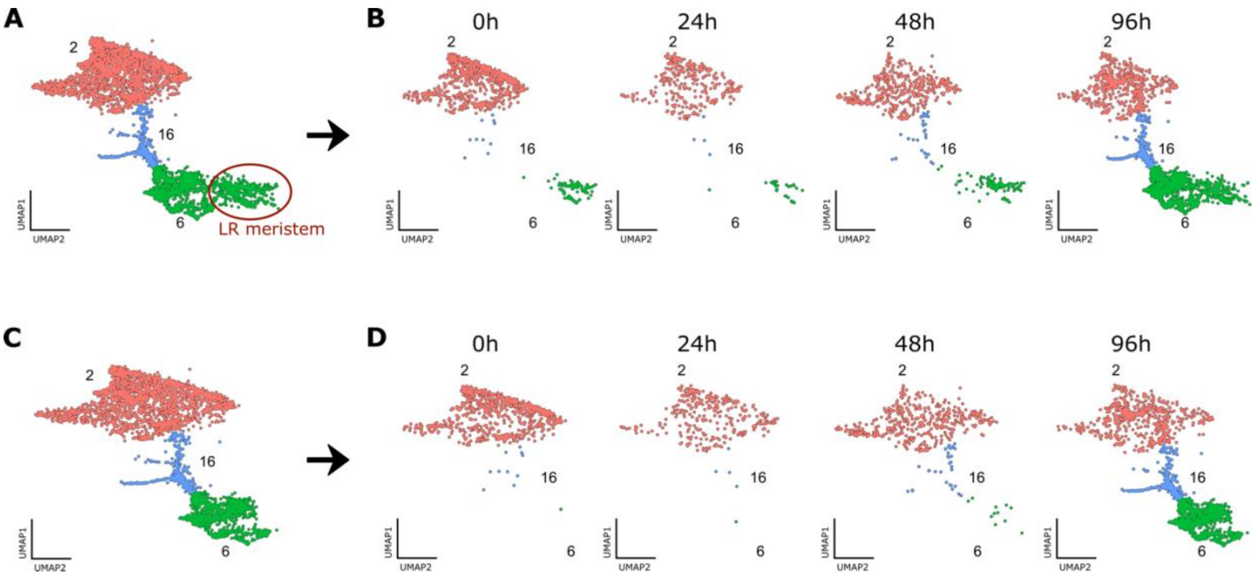


**Fig. S14. Before and after the removal of cells from lateral root (LR) meristem that were mixed on the nodule cluster (cluster 6).** Before the reclustering of the cortex and nodule cells, it was noticed that a subset of cells from cluster 6 was already present in time 0 hpi (A), in opposition to the other cells from this cluster which only appeared after the infection (B). Investigating those cells, we detected the expression of marker genes previously described to be present in both nodule meristem and lateral root meristem, such as the *PLETHORAS* genes. Since the nodule meristem cells are not present at 0 hpi, we inferred that those cells are from the lateral root meristem. This conclusion is supported by the proximity of those cells to cluster 15 in the UMAP space, indicating a similarity in their expression profile. Cluster 15 was annotated as lateral root. To avoid spurious results due to the presence of those cells, they were removed from the dataset before the reclustering (C and D).

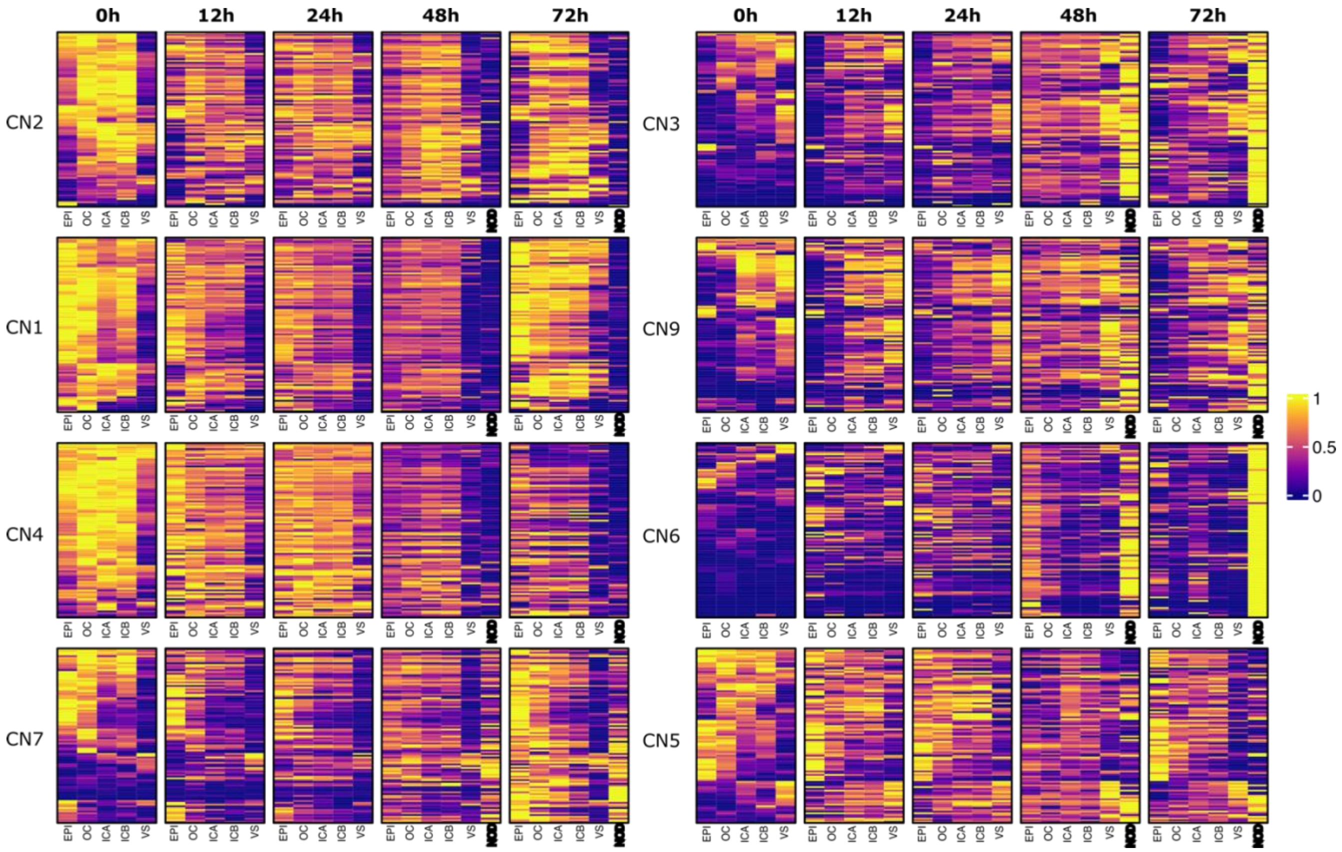


**Fig. S15. Expression profile of the 100 most specifically expressed genes of the 8 subcluster obtained after reclustering cells from cortex and nodule in the LCM dataset.** The subcluster are show in the order they appear in the UMAP plot presented on Figure 4-A (CN2, CN1, CN4, CN7, CN3, CN8, CN6 and CN5. Each column of the heatmap represents the average of three experimental replicates in the LCM dataset (see Methods). The expression was scaled by row. EPI = Epidermal cells, OC = Outer cortical cells, ICA = Inner cortical cells at the xylem poles, ICB = Inner cortical cells between the xylem poles, VS = vasculature, NOD = Nodule. Note that Nodule cells were only sampled at 48 hpi and 72 hpi.

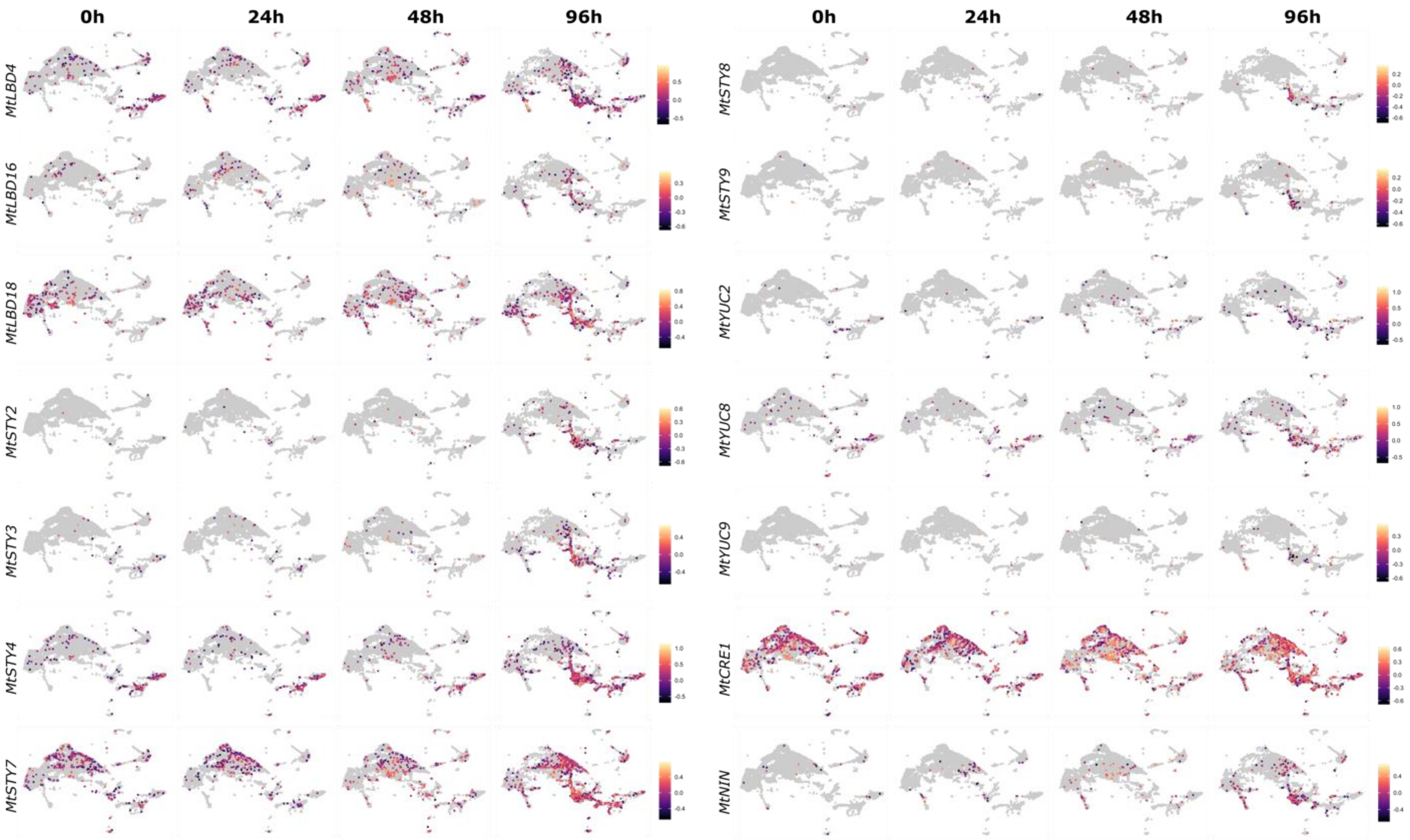


**Fig. S16. Expression profile of all annotated LBD, STY and YUCCA genes in the *M. truncatula* genome (v.5, release 1.9) according with the timepoint after the infection. The expression of *MtCRE1* and *MtNIN*, key regulatory elements of RNS, is show for comparison.** The UMAP plots represents the combination of the cells captured in both Jemalong A17 and *sunn-4*. In each UMAP plot, it is shown the log10 of the expression, which is the default in Monocle3.
